# Self-organizing physical and biochemical interactions explain diverse behaviours in *Physarum polycephalum*

**DOI:** 10.64898/2026.05.07.723662

**Authors:** Linnéa Gyllingberg, Abid Haque, Subash K. Ray, Gregory Weber, Jason M. Graham, Simon Garnier

**Affiliations:** Department of Mathematics, University of California, Los Angeles, CA, USA; Federated Department of Biological Sciences, New Jersey Institute of Technology, Newark, NJ, USA; Federated Department of Biological Sciences, Rutgers University-Newark, Newark, NJ, USA; Department of Biology, University of Indianapolis, Indianapolis, IN, USA; Department of Mathematics, University of Scranton, Scranton, PA, USA

## Abstract

How can simple organisms lacking nervous systems encode and transmit environmental signals to generate complex, adaptive behaviours? Using the unicellular organism *Physarum polycephalum* as a model, we identify a unifying mechanochemical mechanism that links intracellular calcium oscillations to large-scale behavioural coordination. We first demonstrate experimentally that local perturbation of the actomyosin cortex is sufficient to induce symmetry breaking and directed migration, even in the absence of nutrient cues. Building on evidence linking calcium concentration to actin depolymerization and contractile relaxation, we develop a mechanochemical tubule model in which self-sustained calcium oscillations are coupled to pressure-driven mechanics. We show that environmental cues, encoded through the local modulation of these oscillations, give rise to directed transport and the redistribution of biomass. By extending this framework to a two-dimensional phase-field model, we demonstrate that this mechanism is sufficient to generate a diverse set of slime mould behaviours, including chemotaxis, network formation, and balancing exploration–exploitation trade-offs. In doing so, we provide a single mechanistic framework linking intracellular dynamics to organism-scale behaviour across spatial and temporal scales. Our work shows that these sophisticated behaviours can emerge from the modulation of self-sustained oscillations coupled by diffusion, providing a physically grounded mechanism for information processing in non-neural organisms and offering insight into the evolutionary origins of coordinated behaviour.

## Introduction

Living systems have developed remarkable capacities to process information, make decisions, and adapt to changing environments. While most research on this topic has focused on neural organisms– that is, organisms whose behaviour is controlled by a nervous system–growing evidence show that these capabilities are also found in non-neural ones, such as unicellulars [1, 2, 3]. This raises a fundamental question: how can information be encoded, transmitted, and integrated in such organisms in the absence of a dedicated information processing apparatus [4]? Existing studies suggest that self-sustained oscillations–for instance, cyclical calcium dynamics–may encode and transmit various kinds of information through their amplitude, frequency, and phase [5, 6, 7]. However, the link between these intracellular oscillatory dynamics and the macroscopic functional behaviour of the organism is still largely unexplored. In other words, how do molecular mechanisms translate into adaptive outcome at the organism scale?

A natural test bed to address this question is the acellular slime mould, *Physarum polycephalum*. Despite being a single multinucleate cell, it can span over hundreds of square centimetres and move at speeds up to 5 cm h^−1^ [4]. At the organism scale, *Physarum* exhibits a wide range of adaptive behaviours, including directed migration [8], forming efficient transport networks between food sources [9], and balancing exploration-exploitation trade-offs [10]. Its cytoplasm consists of a contractile outer layer surrounding a fluid interior that transports nutrients and organelles [11]. Periodic contractions of the tubules are driven by the actomyosin cortex, generating mechanical forces that induce oscillatory cytoplasmic flow, known as shuttle streaming [12]. These contractions give rise to dynamics across multiple timescales, from minute-scale oscillations to hour-scale network reorganisation.

The membrane contractions are closely linked to intracellular calcium oscillations, which can persist even in cell homogenates, indicating that the underlying biochemical dynamics are self-sustained and can operate independently of the mechanical system [13, 14, 15, 16]. Moreover, calcium signalling has been shown to influence behavioural differences in *Physarum*, including variation in exploration strategies and responses to food sources [17]. Calcium signalling, including intracellular calcium oscillations, is not unique to *Physarum* but is highly conserved across eukaryotic life. It serves as a fundamental intracellular communication mechanism, suggesting that calcium-based dynamics may represent an evolutionarily ancient substrate for coordinating activity within living systems [18, 19, 20]. In *Physarum*, this makes calcium a strong plausible candidate for mediating the translation of local environmental inputs into organism-scale behaviour, enabling information about local conditions to be encoded, propagated, and integrated across the cell through self-sustained dynamics. Using a computational approach, we show here how this putative mechanism can explain the emergence of several well-documented functional behaviours in *Physarum*.

While a wide range of models have been developed to explain different aspects of *Physarum* behaviour, these approaches typically operate at distinct spatio-temporal scales. At short time scales, mechanochemical models describe intracellular oscillations and contraction dynamics [21, 22]. At intermediate time scales, transport models explain shuttle streaming and the propagation of signals across the network through peristaltic flow and advection-mediated feedback [12, 23]. At longer time scales, models of network formation and path optimisation capture morphological adaptation through reinforcement mechanisms [24, 25], while treating internal dynamics phenomenologically. Extensions of these frameworks incorporate oscillatory dynamics into network formation [26], and recent frameworks are beginning to couple oscillatory transport and morphology across scales [27]. Nevertheless, in these recent extensions oscillations are imposed rather than derived from an autonomous biochemical oscillator and therefore do not explain how intracellular dynamics arise from underlying biophysical processes or how they encode and propagate environmental information. More broadly, despite recent advances, no existing model unifies these scales within a single mechanistic framework or captures how local environmental information is integrated through intracellular oscillations to generate coordinated behaviour at the organism scale.

To address this gap, we combine experiments with mathematical models to identify the mechanism linking intracellular signalling to organism-scale behaviour in *Physarum*. We first test whether local mechanical asymmetries are sufficient to drive directional behaviour by perturbing the cortical actin network. Using localized disruption of actin polymerization, we show that weakening the cortex on one side of the cell is sufficient to induce symmetry breaking and directed biomass redistribution, even in the absence of a nutrient cue. This strongly supports the hypothesis that directional behaviour can arise from local modulation of mechanical properties alone. We then investigate how external stimuli can influence behaviour by modulating intracellular processes that regulate the cortical cytoskeleton. Given extensive evidence linking calcium dynamics to actin depolymerization and contractile relaxation [28, 29, 30, 31, 32], we propose that self-sustained calcium oscillations provide the underlying signalling mechanism that generates and coordinates these mechanical asymmetries. To test this, we develop a mechanochemical modelling framework that couples a self-sustained intracellular calcium oscillator to the contractility of the cell’s membrane and the resulting pressure-driven mechanical dynamics. We first formulate a one-dimensional model of a contractile tube, and then extend this approach to two spatial dimensions using a phase-field description of the entire cell. Within this unified framework, external stimuli locally modulate calcium dynamics, and this modulation propagates through diffusive coupling to generate spatially structured patterns of mechanical activity, leading to symmetry breaking, directed transport, and large-scale morphological reorganization. This allows us to account for a range of observed behaviours, including directed migration, network formation, and exploration–exploitation trade-offs, using a single underlying mechanism. More broadly, our results show how diffusive intracellular and oscillatory dynamics, when coupled to mechanical processes, can generate organism-wide functional behaviours, providing insight into how adaptive coordinated action may arise from ancient, non-neural signalling systems.

## Results

### Experimental results

To identify the cellular mechanism underlying directional behaviour in *Physarum polycephalum*, we designed an experimental setup that directly perturbs cortical actin at defined spatial locations. Isolated *Physarum* tubules were exposed to symmetric and asymmetric nutrient cues, or to localized application of the actin-depolymerizing agent cytochalasin-D. Across all conditions, we measured actin abundance at four positions along the tubule and, in asymmetric cases, recorded migratory outcomes (Fig. 1).

**Fig. 1.**
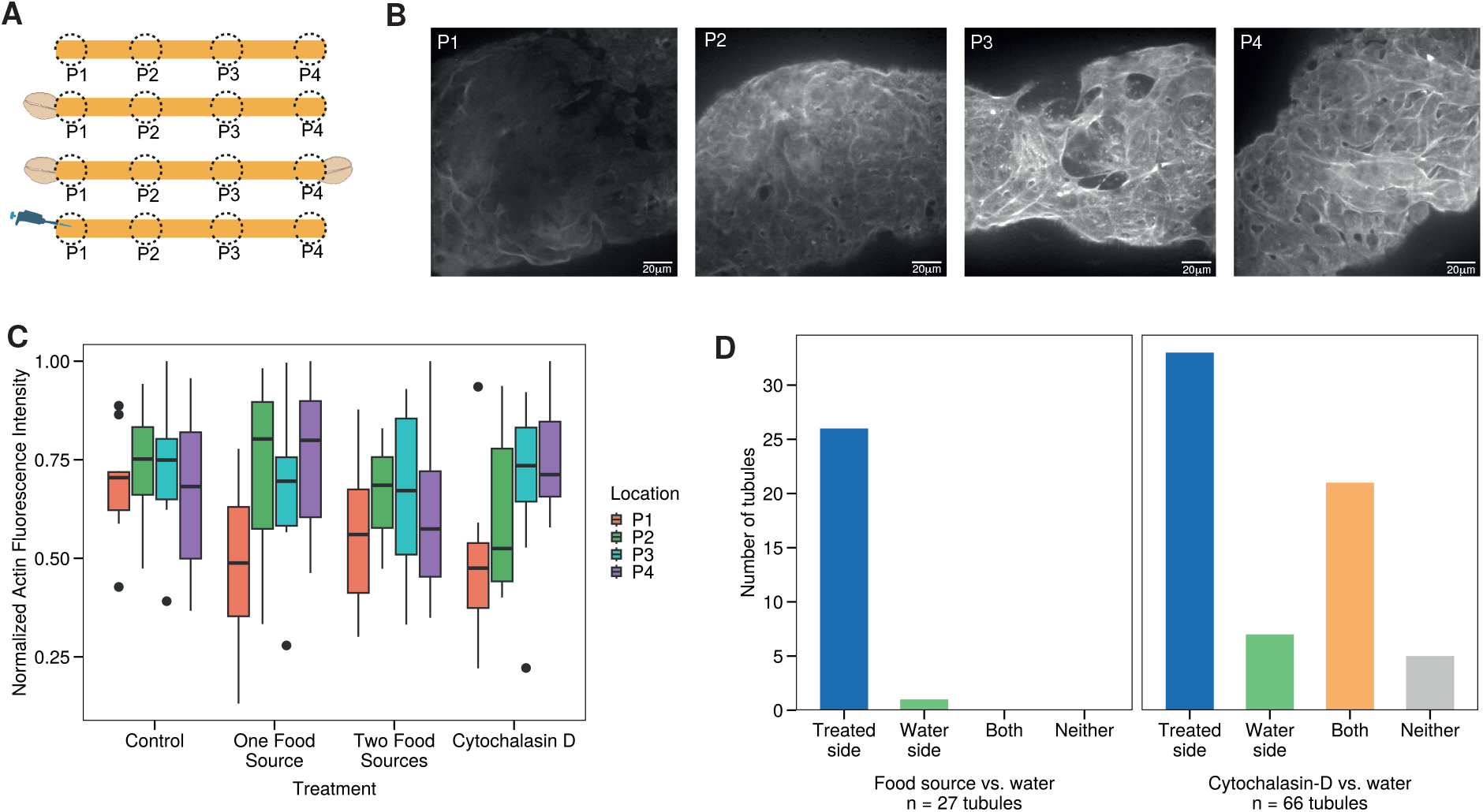
Experimental results. (A) Schematic figure illustrating the four choice conditions used in the migration experiments and actin imaging. The illustration represents, from top to bottom, control condition (no food), asymmetric food condition (food on left side only), symmetric food condition (food on both sides), and the asymmetric cytochalasin-D choice condition. All four conditions were imaged for actin flourescence from four equidistant locations (P1, P2, P3, and P4). However, only the asymmetric conditions were used when studying migration outcomes. (B) Example of the normalized maximum intensity projections of the actin cell cortex at the four equidistant location along the tubule, when exposed to a food source at point P1. (C) Distribution of normalized mean cortical F-actin fluorescence intensity measured at four equidistant positions (P1–P4) along *Physarum* tubules across experimental conditions (control, one food source, two food sources, and localized cytochalasin-D). Values are normalized within each tubule. Under the asymmetric conditions (one food source and localized cytochalasin-D) actin abundance is reduced at the perturbed site (P1) relative to more distal sites. In the symmetric food condition (two food sources), actin abundance is slightly lower at both ends (P1 and P4), whereas in the control condition, no consistent spatial structure is observed. (D) Distribution of final migration outcomes under asymmetric conditions. Left: when presented with a food source versus a water-soaked non-nutrient agar block, tubules migrated toward the food source in 26/27 cases. Right: when presented with cytochalasin-D versus water-soaked non-nutrient agar block, tubules migrated toward the treated side in 33/66 cases, toward the control in 7/66, toward both in 21/66, and toward neither in 5/66.

When presented with an asymmetric nutrient source, initially unpolarized tubules exhibited a significant preference (proportionality test, *p <* 10^−5^, *N* = 27) for the more nutrient-rich food source on the left side, with a large number (26 out of 27) of tubules rejecting the non-nutrient side, and 1 out of 27 tubules retracting from both the food source and the agar-water block **(Fig. 1D)**. This transition was accompanied by a pronounced spatial gradient in cortical F-actin abundance, where the F-actin density was reduced at the leading edge relative to distal regions **(Fig. 1)**. In this condition, with one food source on the left side of the tubule and a non-nutrient agar block soaked in water (*N* = 13), we found that the actin fluorescence intensity at the location closest to the food source (P1) was significantly lower than the fluorescence intensity observed at other locations P2 (*p <* 0.001), P3 (*p <* 0.01), and P4 (*p <* 0.001). Directed migration thus emerges alongside a local weakening of the cortical actin network, suggesting that spatial modulation of cortical mechanics may bias intracellular mass redistribution.

To test whether such local mechanical asymmetries are sufficient to drive directional behaviour, we applied cytochalasin-D locally to one side of the tubule (*N* = 10). This perturbation reproduced the same qualitative actin gradients observed near nutrient sources, and we found that the actin fluorescence intensity at the location closest to the food source (P1) was significantly lower than the fluorescence intensity observed at other locations P3 (*p <* 0.05) and P4 (*p <* 0.01). Crucially, this was sufficient to induce a significant preference (proportionality test, *p <* 0.01, *N* = 66) of directed biomass redistribution toward the perturbed side, even in the absence of a nutritional cue, with 33 out of 66 tubules exclusively choosing the side with the cytochalasin-D agar block, while 7 out of 66 chose only the agar-water block (Fig. 1 C and D).

Consistently across asymmetric conditions, regions exhibiting reduced cortical actin were preferentially reinforced and became sites of net biomass accumulation. Taken together, these results support the hypothesis that localized actin depolymerization lowers cortical tension and hydraulic resistance, thereby generating mechanically driven intracellular asymmetries that bias migration direction.

### Tubule model

While these experiments show that directional behaviour arises from local modulation of cortical mechanics, they do not explain how environmental information is encoded into mechanical perturbations. Experimental studies have shown that calcium regulates actin depolymerization and contractile relaxation, and that chemical stimuli can modulate intracellular calcium levels through pathways such as IP_3_ signalling [14, 33]. We therefore hypothesize that self-sustained calcium oscillations provide the signalling layer that links external inputs to cortical mechanics (Fig. 2) .

**Fig. 2.**
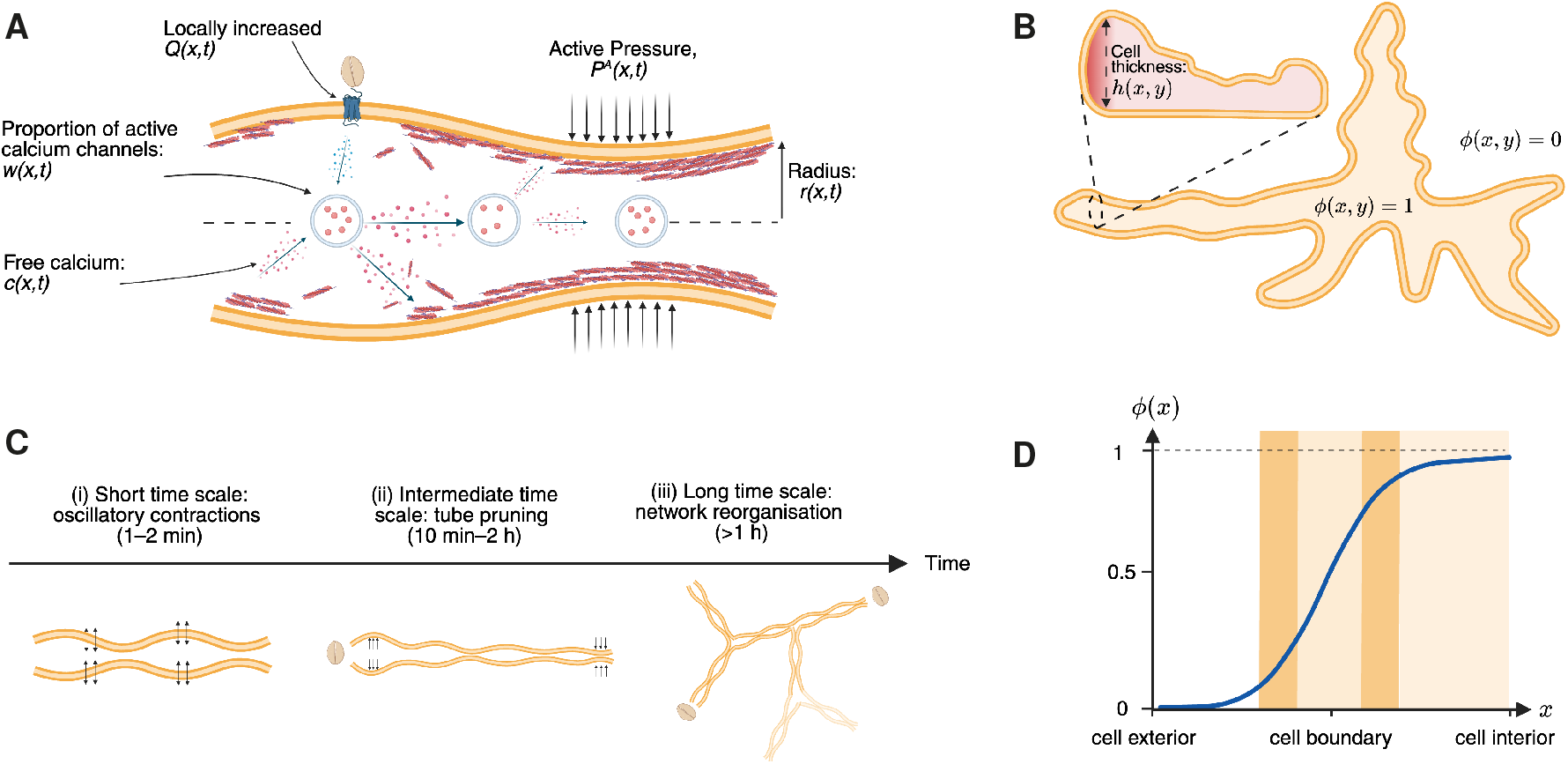
Schematic overview of the mechanochemical framework linking intracellular calcium dynamics to organism-scale behaviour in *Physarum polycephalum*. (A) Local environmental stimuli, such as nutrients, increase the permeability of IP_3_-sensitive calcium channels, represented by *Q*(*x, t*), leading to elevated cytosolic calcium *c*(*x, t*). Calcium-induced calcium release (CICR) generates a positive feedback effect, while re-sequestration by pumps provides negative feedback effect, resulting in self-sustained calcium oscillations. Elevated calcium promotes actin depolymerization, locally reducing cortical tension (*P*^*A*^(*x, t*)) and increasing tube radius (*r*(*x, t*)), while low calcium levels restore actin polymerization and contractility. Calcium further spreads through diffusion, coupling neighbouring regions. (B) Phase-field representation of the cell, where *ϕ*(*x, y*) = 1 denotes the inside of the cell and *ϕ*(*x, y*) = 0 the outside of the cell, and *h*(*x, y*) describes the local cell thickness. (C) Dynamics across time scales: short-time oscillatory contractions (1–2 min), intermediate-time tube pruning and directional biomass redistribution (10 min–2 h), and long-time network reorganization (*>* 1h). The visual organization of panel C was adapted from [34] (CC BY 4.0). (D) Diffuse interface profile of the phase-field variable *ϕ*(*x*) across the cell boundary: the transition between the cell interior and exterior is represented as a sharp, but smooth transition.

Based on this hypothesis, we develop a mechanochemical framework in which intracellular calcium oscillations regulate cortical contractility and thereby generate active pressure. Local modulation of calcium dynamics leads to spatial variations in contractile stress, producing pressure gradients that drive fluid flow and deformation of the cell.

To this end, we start by formulating a one-dimensional model of a thin *Physarum* tubule of length *L* = 1 cm and with a mean radius of 100 µm. The tubule is defined on a spatial domain *x* ∈ [0, *L*], and modelled as a contractile active fluid in which calcium dynamics modulate actomyosin contractility and drive flow through pressure gradients. The model captures four coupled processes: (1) cytoplasmic calcium oscillations, (2) calcium-dependent regulation of cortical contractile pressure, (3) mechanical feedback through changes in tube radius and fluid flow, and (4) modulation of intracellular calcium by external stimuli.

The self-sustained calcium oscillations are described by a spatial extension of the model proposed by Poledna et al. [35], in which the calcium concentration *c*(*x, t*) and the fraction of calcium channels *w*(*x, t*) available for calcium release regulate each other:

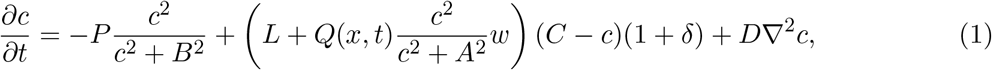

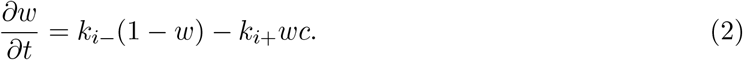

The first term in Eq. (1) represents calcium removal by pumps, while the second term describes calcium release through IP_3_-sensitive channels. This release increases with both the local calcium concentration and the fraction of channels available for calcium release, representing calcium-induced calcium release (CICR), which provides a positive feedback mechanism. In contrast, calcium removal by pumps acts as a negative feedback, restoring basal calcium levels. The interplay between these two processes generates self-sustained oscillations in calcium concentration. We extend Poledna et al’s model by including a diffusion term, accounting for the diffusive transportation of calcium at physiological concentrations [18].

A key parameter in this equation is *Q*(*x, t*), which represents the effective permeability of calcium channels and encodes local activation of IP_3_ receptors. In the model, *Q*(*x, t*) serves as the interface between external stimuli and intracellular dynamics. To capture intrinsic variability in signalling and environmental sensing, we allow *Q*(*x, t*) to exhibit small stochastic fluctuations, described in detail in the Supplementary Section SI2.3. In practice, a locally elevated *Q*(*x, t*) should be interpreted as a higher local concentration of an attractant, such as a nutrient source.

To couple biochemical oscillations to mechanics, we model the local calcium concentration, *c*(*x, t*) as a variable controlling the local pressure, consistent with experiments showing that calcium oscillations precede and drive contractions in *Physarum* [14, 36]. Adapting the framework of Teplov et al. [21], we write:

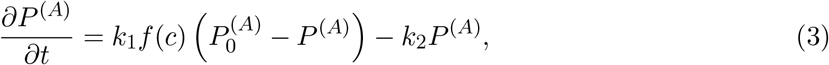

where *f* (*c*) is a monotonically decreasing sigmoidal function of the calcium concentration (see Methods). The local differences in pressure give rise to pressure gradients, deforming the local radius of the tubule, *r*(*x, t*):

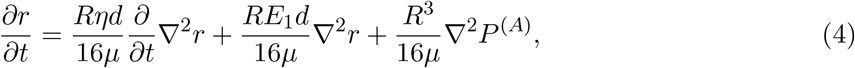

The first term gives viscoelastic relaxation of the cortex, the second term represents elasticity, and the third term captures active deformation driven by pressure gradients. In summary, the spatiotemporal changes in radius are driven by active pressure, setting up pressure gradients and fluid flow, which in turn, are driven by the underlying biochemical oscillations. The full derivation of the model as well as exact parameters are found in the Supplementary Section SI2.

We first study the system in the absence of external stimuli, where the mean value of *Q*(*x, t*) is spatially homogeneous across the tubule. For a homogeneous value *Q* = 0.3, the model exhibit spatially synchronous, self-sustained oscillations in calcium concentration, active pressure, and tubule radius, with a period of approximately 100 seconds (Fig. 3A), consistent with experimentally observed calcium and radius contractile oscillations, which typically have periods around 1-3 minutes [37, 15]. Fig. 3A shows a representative time series at the midpoint of the tube (*x* = 0.5).

**Fig. 3.**
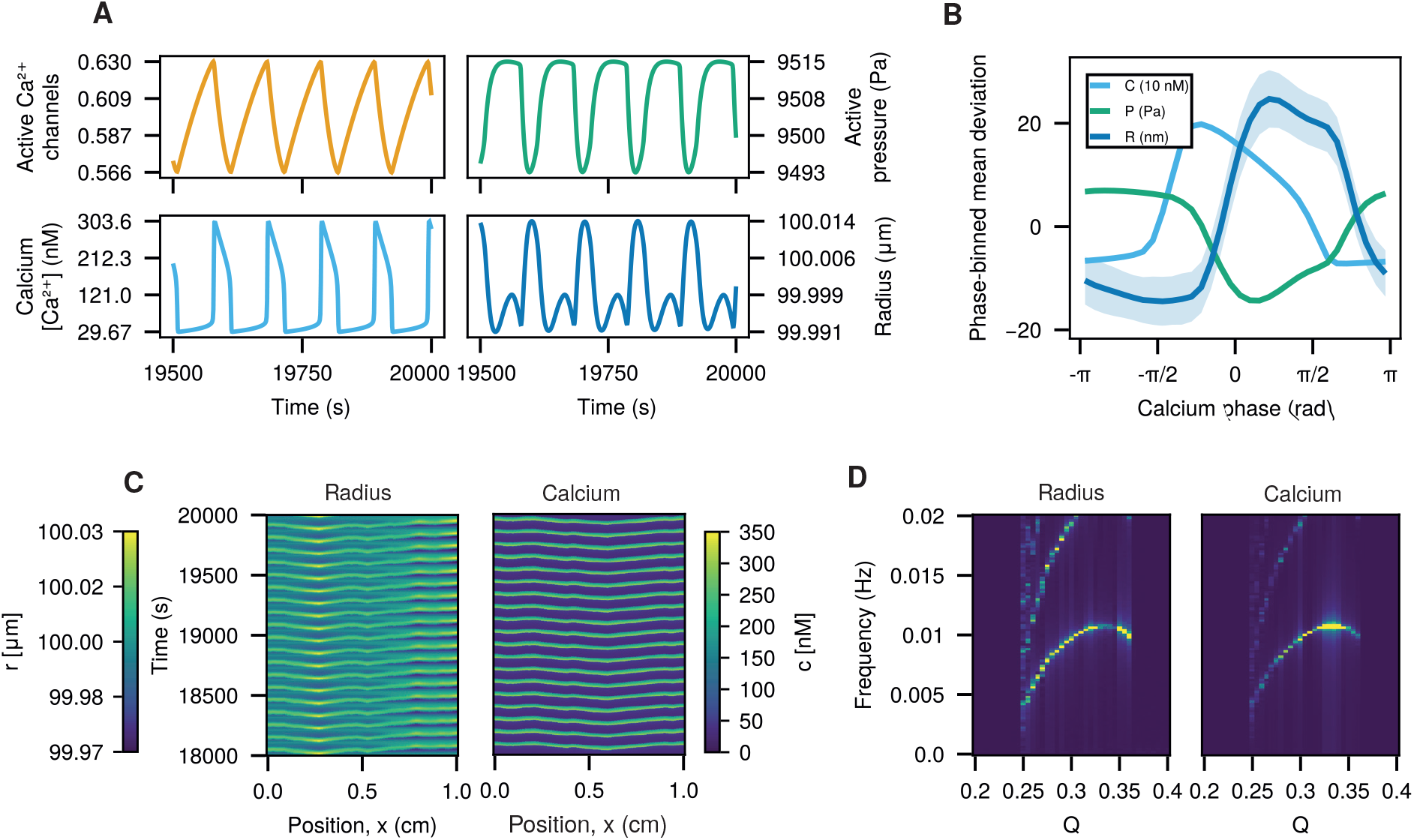
Baseline oscillatory dynamics in the one dimensional mechanochemical tubule model under spatially homogeneous *Q*(*x, t*). (A) Representative time series at the tube mid point, *x* = 0.5, after 19500 seconds, showing self-sustained oscillations in proportion of active calcium channels, calcium concentration, active pressure and radius, with a frequency around 0.01 Hz. (B) Phase relation between calcium concentration, active pressure, and radius as functions of the calcium oscillation phase at the tube midpoint. Lines show the mean across simulations, and shaded regions indicate one standard deviation. (C) Space-time kymographs of radius and calcium concentration along the tubule, illustrating standing-wave patterns when *Q*(*x, t*) is homogenous. (D) Oscillation frequency of calcium concentration and tubule radius as a function of the calcium channel permeability parameter *Q*, showing the onset and disappearance of oscillations as *Q* is varied.

To understand how the biochemical and mechanical variables are related during a single oscillation cycle, we analyse the phase relationship between calcium concentration, active pressure, and radius at the midpoint of the tube. Calcium and radius oscillate approximately in antiphase, with active pressure peaking roughly *π* after the calcium maximum (Fig. 3B). These phase relations are consistent with experimental observations in *Physarum* [14, 36].

The system exhibits standing-wave patterns with synchronous calcium oscillations across the tubule (Fig. 3C). To examine how environmental modulation affects these dynamics, we vary the calcium channel permeability *Q*. Increasing *Q* leads to the onset of oscillations beyond a critical threshold (*Q* ≈ 0.25). As *Q* is increased further, the oscillation frequency increases substantially over an extended range of *Q*, followed by a plateau, and then exhibits a slight decrease before oscillations are lost at higher values (*Q* ≳ 0.37). This is illustrated in the bifurcation diagram in Figure 3D. These findings are consistent with experimental observations in *Physarum*, where increasing attractant concentration leads to increased oscillation frequencies [38]. Motivated by the sustained calcium oscillations observed in *Physarum*, we focus on the parameter regime exhibiting persistent oscillatory dynamics.

While the model with a spatially homogenous *Q* value describes the short time scale coordination of the biochemical and mechanical variables (Fig. 2C (i)), it does not capture the intermediate time scale of biomass redistribution and symmetry breaking that occur when environmental cues vary across space (Fig. 2C (ii)). We therefore consider the case where *Q* is spatially inhomogeneous.

In Fig. 4A,C we simulate the introduction of a food source at the left end of the tubule at time *t* = 20000, implemented as *Q*(*x, t*) = 0.35 for 0 ≤ *x* ≤ 0.1 and *t >* 20000. This local increase in *Q* alters the oscillation pattern of the radius of tubule as well as the calcium concentration in the tubule (Fig.4A). The elevated channel permeability near the food source produces larger calcium transients in that region, which in turn generate a spatial gradient in active pressure. As a consequence, the baseline standing-wave regime transitions into travelling waves that propagate toward the food source. This transition is accompanied by a modulation of contraction amplitude that originates at the stimulated end and spreads across the tubule. These kymographs are consistent with experimental observations by Alim et al. [23], who reported that local stimulation induces propagating changes in contraction amplitude that spread from the site of stimulation across the tubule network of *Physarum*.

**Fig. 4.**
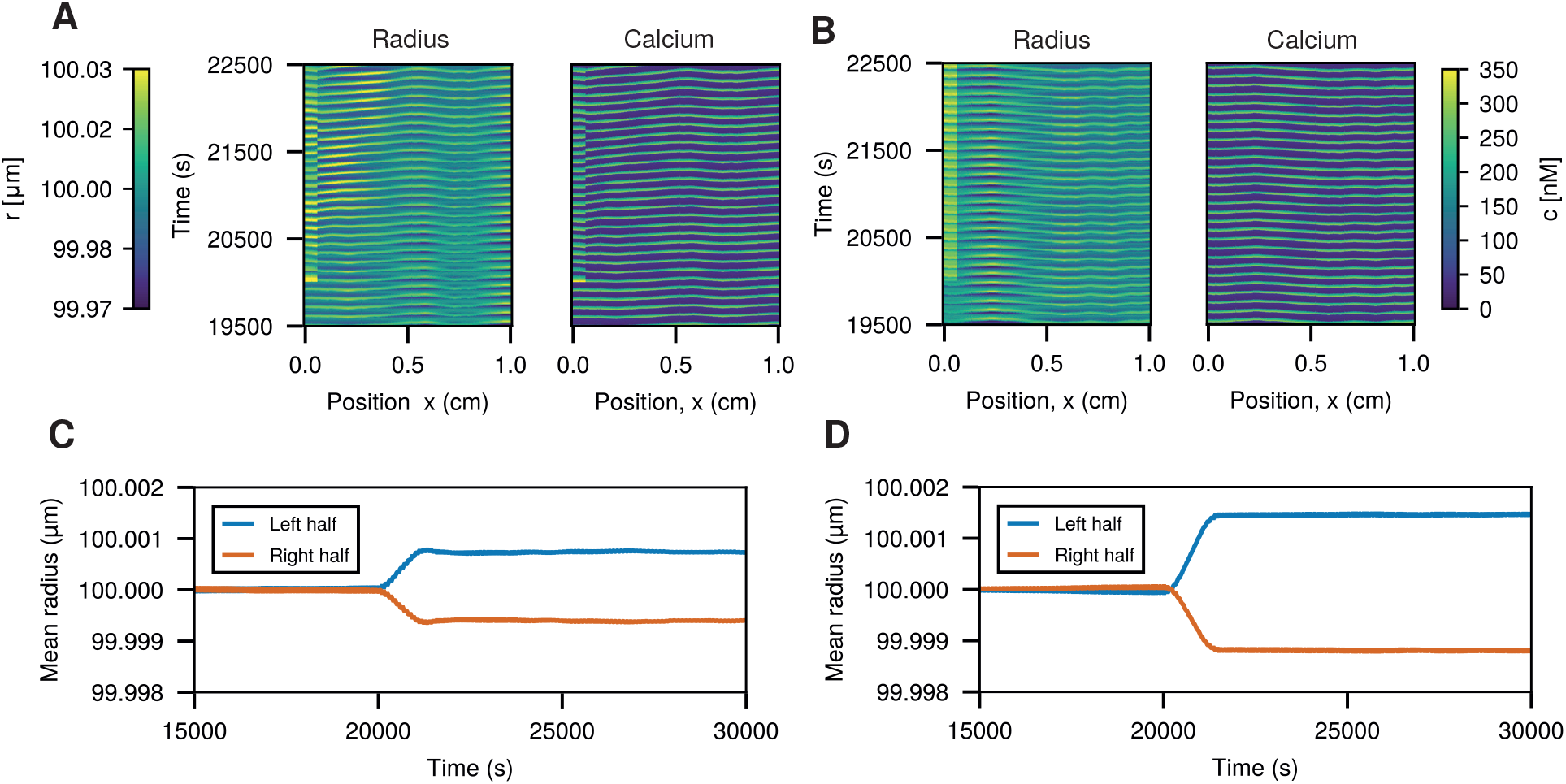
Effects of local perturbations in the one-dimensional tubule model. (A,B) Kymographs of the tubule radius and calcium concentration after the introduction of a local perturbation at the left end of the domain at *t* = 20,000. In (A), a local increase in *Q* to 0.35 from the baseline value 0.3 (modelling the presence of a food source) alters the oscillatory pattern, producing travelling waves that originate at the stimulated end and propagate across the domain. In (B), a local reduction in the actin polymerization forward rate, *k*_1_, modelling the presence of Cytochalasin-D, causes the tube to thicken on the left-hand side, while leaving the oscillatory pattern for the radius largely unchanged with no consistent wave propagation direction. The calcium dynamics are unaffected. (C,D) Time evolution of the mean radius in the left and right halves of the tubule for the two perturbations. In both cases, after stimulation at time *t* = 20,000, the perturbed side thickens, whereas the unaffected side becomes thinner. Taken together, these panels show that spatially inhomogeneous *Q* induces travelling waves together with mass redistribution, whereas reducing *k*_1_ leads to mass redistribution without directed wave propagation.

This transition is accompanied by symmetry breaking in tube radius: the stimulated side thickens while the opposite side thins, leading to a persistent redistribution of biomass. We quantify this by comparing the mean radius of the left and right halves of the tubule, which are approximately the same prior to stimulation but evolve toward a stable asymmetric state, in which the stimulated side is thicker and the opposite side thinner (Fig. 3C).

To assess robustness across stochastic realizations, we performed 30 independent simulations. Following stimulation, wave propagation becomes consistently biased toward the high-*Q* region. This directional bias is accompanied by a redistribution of biomass toward the stimulated side, increased calcium asymmetry, and a shift in the spatial structure of the oscillatory pattern. All effects are statistically significant (see Supplementary Fig. SI4).

To replicate the cytochalasin experiments, we locally reduce the actin polymerization rate *k*_1_, which in the model corresponds to cytochalasin-D disrupting actin polymerization. Specifically, we decrease *k*_1_ by 2% at the left end of the tubule at time *t* = 20 000 (Fig. 4B,D).

In contrast to the asymmetric *Q* case, the spatiotemporal wave pattern remains unchanged, with no systematic shift in wave direction. Instead, lowering *k*_1_ locally reduces active pressure, creating a static pressure difference along the tubule without inducing travelling waves. Nevertheless, the system undergoes symmetry breaking in tube radius: the perturbed side thickens while the opposite side thins, leading to a persistent volume imbalance without directed wave propagation.

Again, these finding are consistent across 30 stochastic realizations; we observed no significant change in wave direction or spatial mode, while biomass asymmetry increases significantly. This indicates that local weakening of actin polymerization redistributes mass without inducing travelling waves (see Supplementary Fig. SI5).

### Phase field model

The one-dimensional model captures several key aspects of *Physarum* dynamics observed in experiments at the tubule and biomass-redistribution scales, corresponding to spatiotemporal scales (i) and (ii) in Fig. 2C. These include self-sustained oscillations, the experimentally observed antiphase relationship between calcium concentration and tubule radius, as well as symmetry breaking and redistribution of biomass when food is present. However, since the model represents the organism as a single tubule, it captures only the biomass imbalances that precede movement and cannot describe whole-cell migration or large-scale morphological rearrangements, corresponding to spatiotemporal scale (iii) in Fig. 2C. We therefore extend the model to two spatial dimensions by incorporating the calcium-driven contractile mechanism into a phase-field description [39, 40] of the plasmodium, allowing intracellular signalling to regulate cell shape, thickness, and movement directly.

In this framework, the plasmodium is represented by a scalar field *ϕ*(*x, y, t*) ∈ [0, 1], which describes the local presence of cytoplasm (Fig 2B). Regions with *ϕ* ≈ 1 are inside the cell and regions with *ϕ* ≈ 0 are outside the cell, with the cell boundary represented as a thin transition region between these states, rather than a sharp interface (Fig 2 D).

To describe whole-cell deformation, we model the evolution of the interface by passive interfacial relaxation (which smooths the cell boundary), volume conservation, and calcium-modulated active boundary forcing:

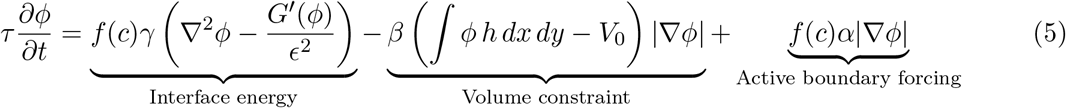

The first term represents passive interfacial tension, implemented through a double-well free-energy formulation commonly used in phase-field models [41, 42, 43]. This term penalizes sharp gradients in *ϕ* and thereby stabilizes the interface. In contrast to classical phase-field formulations, the interfacial stiffness is modulated by the local calcium concentration *c*(*x, y, t*), allowing intracellular biochemical signals to locally soften or stiffen the cell boundary.

The second term enforces conservation of the total cell volume. If the integrated phase field ∫ *ϕ h dx dy* deviates from the initial total volume *V*_0_, this term restores the volume through a force acting at the cell boundary (|∇*ϕ*|).

The third term represents calcium-dependent active forces generated by the cortical actomyosin network, which drive local protrusion and retraction of the boundary.

Although the model is defined on a two-dimensional domain, variations in biomass are captured through a thickness field *h*(*x, y, t*), which represents the local height of the plasmodial sheet. This allows the model to account for mass redistribution within the cell. This gives an effective three dimensional cell representation, while being computationally tractable:

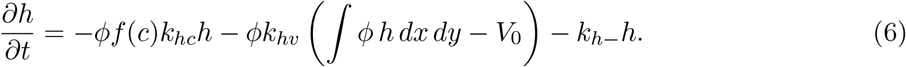

The first term represents calcium-modulated contractility that reduces local thickness. The second term enforces approximate conservation of the total cell volume. The final term represents passive relaxation of thickness in regions where the plasmodium is thinning.

As in the one-dimensional model, intracellular calcium dynamics are governed by the same oscillator coupling the calcium concentration *c*(*x, y, t*) and the fraction of calcium channels available for calcium release *w*(*x, y, t*) (see Methods).

In both the tubule model and the phase field model, we neglect total biomass growth, as movement and redistribution of biomass occur on much shorter time scales (minutes to hours) than cell growth, which in *Physarum* occurs on the order of days [44].

### Baseline behaviour

As a baseline, we first examine how the *Physarum* plasmodium explores space in the absence of food sources, that is, when the calcium channel permeability *Q* is spatially uniform and set to the baseline value 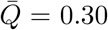. Under these conditions the cell expands approximately isotropically, maintaining an almost circular morphology while exhibiting negligible displacement of its center of mass.

The cell boundary undergoes periodic oscillations, repeatedly expanding and retracting, rather than expanding monotonically, consistent with experimental observations of alternating periods of expansion, stagnation and retraction of *Physarum* fronts [45]. These oscillations are biased outwards, producing a net expansion with an average outward propagation speed of approximately 0.64 µm s^−1^, while the instantaneous speed oscillates between positive and negative values of order 10-20 µm s^−1^ comparable to experimental measurements [45] (see Supplementary Fig. SI6). As a result, the plasmodium gradually spreads outward until it nearly occupies the entire arena.

During this symmetric expansion, intracellular calcium dynamics remain radially symmetric and organized in a circular oscillatory mode, reflecting the absence of any preferred migration direction (see Supplementary Fig. SI7). This non-polarized baseline state is consistent with experimental observations that concentrically expanding plasmodia exhibit symmetric intracellular patterns prior to the onset of directional migration [46].

### Chemotaxis

We next investigate how external cues perturb this symmetric dynamical state. In natural environments, *Physarum* plasmodia respond to spatial gradients of nutrients and other chemical signals by reorienting their cell and migrating toward favourable regions. Such chemotactic behaviour has been documented in slime moulds for decades [47, 8, 48]. Experimental studies further suggest that directional migration is accompanied by intracellular asymmetries in calcium dynamics, with elevated calcium activity observed on the side of the plasmodium facing an attractant [49].

To examine whether our model reproduces this transition from symmetric exploration to directed migration, we introduce a spatially localized nutrient source–implemented as a region where *Q* is elevated–and study how the initially isotropic plasmodium responds to the resulting chemical gradient. Initially, the cell expands approximately symmetrically. Once it encounters the chemical gradient, calcium levels increase in the region facing the nutrient source (see Supplementary Fig. SI8), consistent with with experimental observations [49]. As a consequence, radial symmetry breaks and the cell polarizes, forming an anterior in the direction of the nutrient field. The simulated plasmodium then migrates toward the nutrient source.

Migration speed in slime moulds can be quantified in several ways, for example using the velocity of the center of mass or the speed of the leading edge. Across 30 independent simulations, the center of mass of the plasmodium moves toward the nutrient source with an average speed of 9.87 µm s^−1^, corresponding to a mean displacement of 0.709 mm. The instantaneous leading edge speed alternates between 0 and 20 µm s^−1^, consistent with experimental measurements [45]. As in the baseline case, the cell moves by alternating between forward and backward motion, but with a slightly net forward motion, leading to migration toward the food source, in line with experimental observations of migrating *Physarum* cells [50].

### Network formation

One of the most well-known behaviours of *Physarum* is its ability to self-organize into transport networks that are often described as optimal [51, 9]. To this end, we simulate a simple network experiment, where food is located in three corners of the arena (implemented as circular areas where *Q* is elevated), and the slime mould is placed as a droplet in the mid position of them. Just as in the baseline case, the slime mould starts expanding with radial symmetry. However, when detecting the three food sources, rather than migrating towards one of the food sources, like in the chemotaxis simulations, it starts rearranging its morphology. First, the cell becomes perforated, having multiple gaps and areas with low thickness. This is followed by selective retention and pruning which leads to the cell forming a sparse network of thicker strands connecting the three food sources. This can be seen in Supplementary Fig. SI6C, where the area of the slime mould fist increases to ≈ 2.0 mm^2^, until it touches all the food sources and starts redistributing its biomass. In a subset of realizations the plasmodium does not stay connected: in 8 out of 30 simulations, after reaching all three food sources, the plasmodium splits into two parts, one that connects two of the food sources, and one surrounding the third food source (see Supplementary Fig. SI9). This type of macroscopic splitting into autonomous subunits has been experimentally observed in *Physarum* [52].

### Exploration-exploitation trade-offs

Another remarkable ability of slime moulds is their capacity to balance exploration–exploitation trade-offs, a feature revealed in experiments inspired by the multi-armed bandit problem [10]. Reid et al. placed slime mould biomass at the base of two arms containing discrete agar sites that differed in overall nutrient reward. The first sites on each arm contained food to ensure initial exploration, while the total amount and spatial distribution of food patches differed between arms, defining a low-quality (LQ) and a high-quality (HQ) arm.

In this context, exploration corresponds to biomass spreading along both arms of the arena, whereas exploitation corresponds to sustained growth along only one of the arms. To test whether our model reproduces this behaviour, we simulate in silico the same experimental setup as in [10] using a thin two-arm arena (0.1 mm × 1 mm cross-section) with patchy food distributions along each arm (Fig. 6(A)). We quantify the resulting behaviour by measuring the difference in the furthest distance reached by the leading edge along the two arms, as well as the proportion of replicates that reached the high quality arm first, as in [10].

**Fig. 5.**
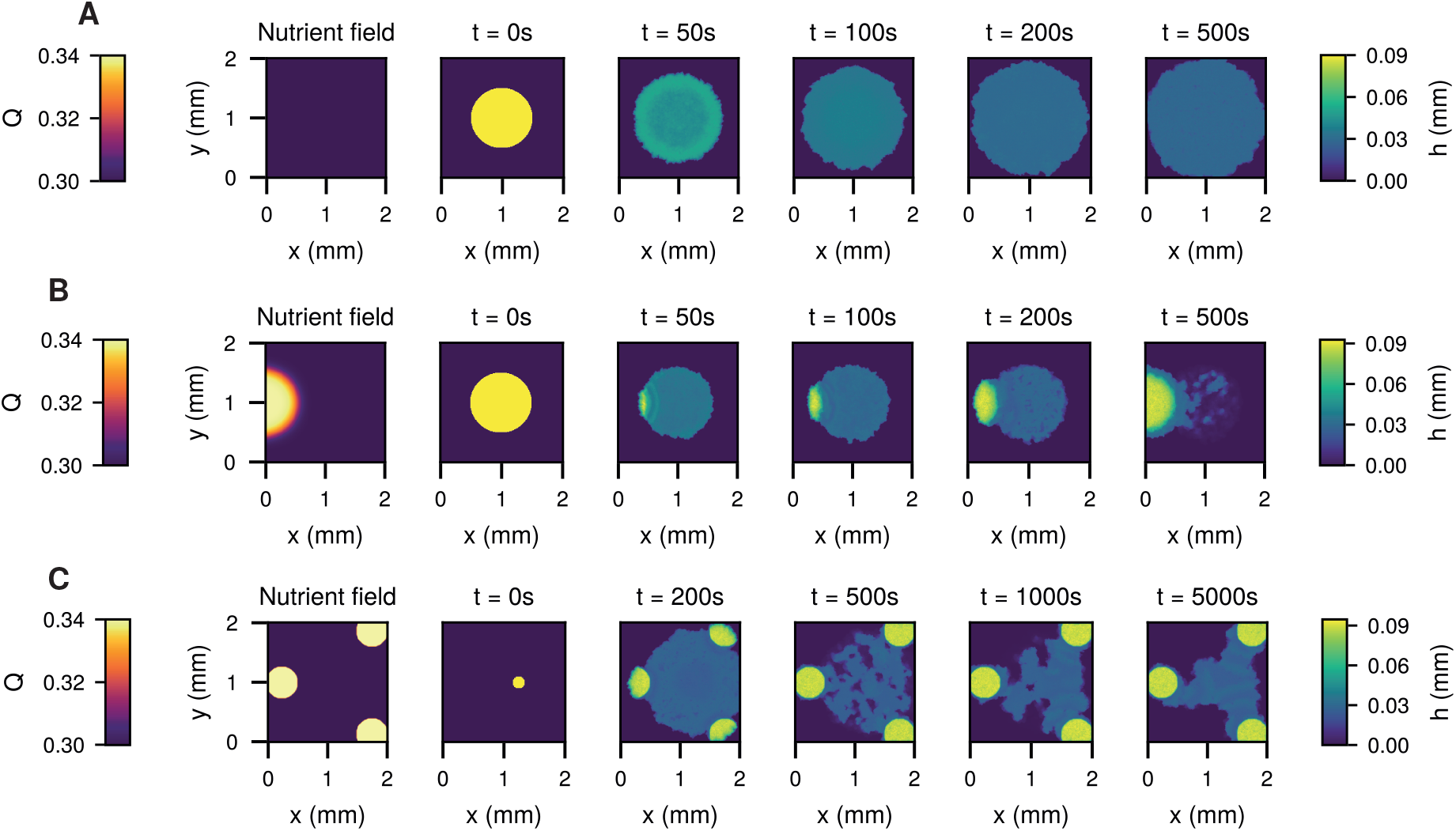
Phase field simulations of shape dynamics under homogeneous and heterogeneous nutrient conditions (A) Baseline setup with a spatially uniform nutrient field, corresponding to constant calcium channel permeability *Q*. Starting from an initially circular configuration, the cell spreads approximately isotropically, with negligible net displacement of its center of mass. (B) Chemotactic setup with a localized nutrient source on the left. When encountering the nutrient gradient, the initially circular cell undergoes symmetry breaking and develops a polarized morphology. It migrates toward the nutrient-rich region. (C) In a heterogeneous nutrient landscape with three localized food sources, the cell starts expanding symmetrically. However, when coming in contact with the three food sources, the cell temporary becomes perforated, having multiple low thickness gaps. This follows by a reorganization of the morphology into thicker strands and eventually these strands prune into a sparse, tree like network connecting nutrient rich regions. Colour indicates local cell thickness *h*(*x, y, t*), and snapshots show the temporal evolution at the indicated times. Panels show a representative realisation – all simulations were repeated 30 times.

**Fig. 6.**
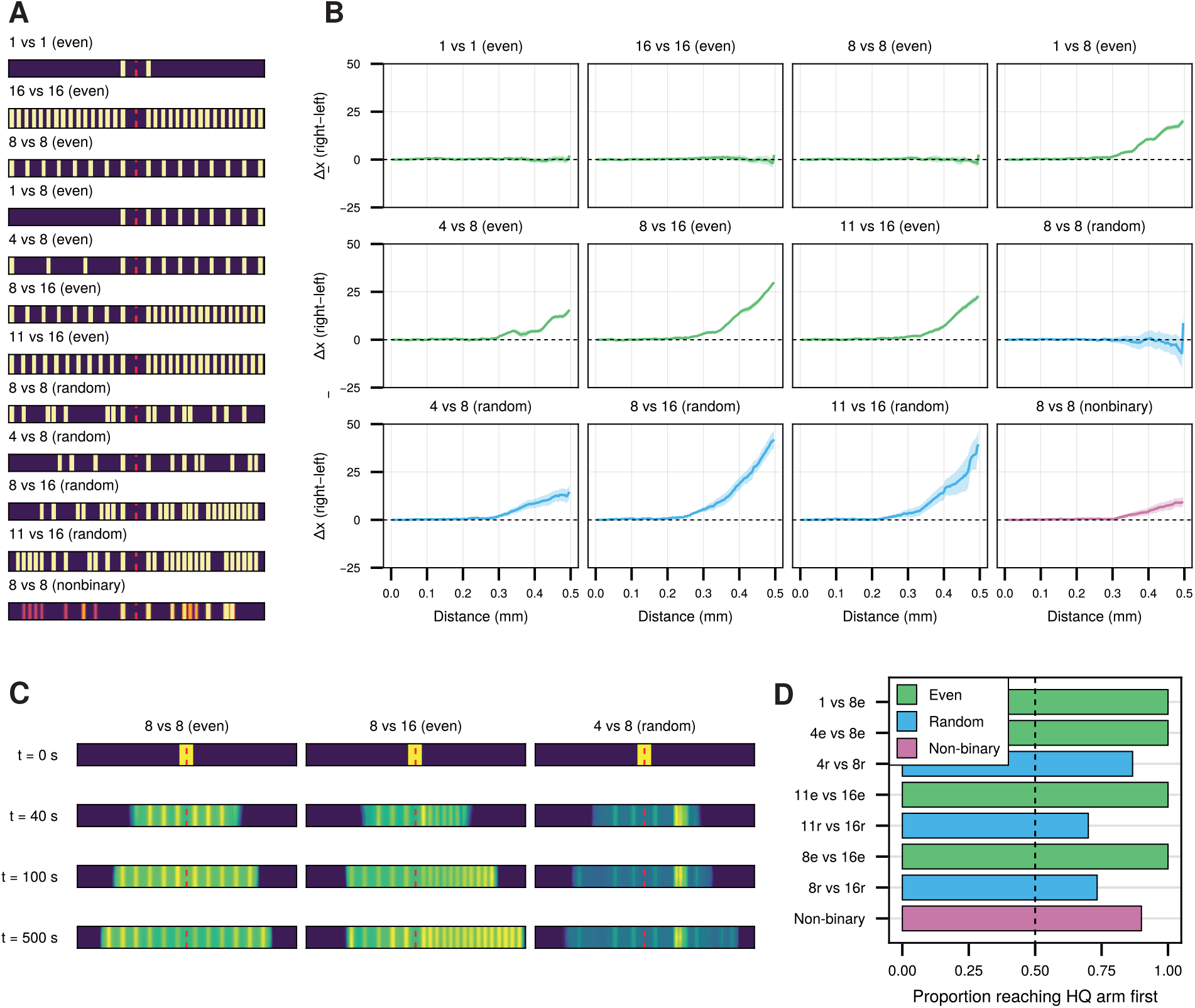
Exploration–exploitation dynamics across experimental conditions. (A) Spatial arrangement of food sources across experimental conditions. The red line indicates the midpoint of the arena. For the random and non-binary conditions, each simulation generates a new arrangement of food patches. (B)Difference in exploration between the high-quality (HQ) and low-quality (LQ) arm across the 12 experimental treatments. The plotted quantity shows the difference in the furthest position reached by the leading edge on the HQ and LQ arms as successive food sites are discovered. Positive values indicate greater exploration of the HQ arm, negative values indicate greater exploration of the LQ arm, and zero indicates equal exploration of both arms. For each treatment, we performed 30 independent simulations; the solid line shows the mean and the shaded region the 95% confidence interval. As in the experimental setup of [10], the first site on each arm was always a food site to ensure initial exploration of both arms. In the random treatments, food patch locations along the arms were randomly generated for each simulation. (C) Representative time points illustrating exploration–exploitation dynamics. In all cases, the cell starts at the midpoint of the arena (*t* = 0). At intermediate times (*t* = 40), both arms are explored equally across all experimental conditions, indicating an exploration phase. At later times (*t* = 100 and *t* = 500), movement becomes biased toward one arm in asymmetric conditions, while remaining symmetric in the 8 vs 8 condition where no preference is present, reflecting the transition from exploration to preferential exploitation in the asymmetric cases. (D) The proportion of simulation that successfully reached the end of the high-quality (HQ) arm before reaching the end of the low-quality (LQ) arm in each of the asymmetric treatments. The dashed line indicates chance level (0.5), with values above this threshold reflecting a bias toward the HQ arm.

For the experimental setups with equally rewarding arms, the in-silico slime mould continues to explore both arms equally, resulting in a near-zero difference between the HQ and LQ arms (Fig. 6B,C). Across the nine treatments with unequal reward, the cell initially explores both arms, but over time the leading edge advances further along the high-quality arm, indicating a shift toward preferential exploitation of the higher-reward environment (Fig. 6B,C). Both the speed of this transition and the magnitude of the bias depend on the spatial distribution of food patches. The proportion of simulations that reach the end of the HQ arm before the LQ arm provides an additional measure of this bias (Fig. 6D). In the evenly distributed unequal-reward treatments, the plasmodium reaches the HQ arm first in all 30 simulations, whereas this preference is weaker in the randomly distributed and non-binary treatments. Together, these results agree well with the experimental findings of Reid et al. [10]. Our model shows that this bias emerges from nutrient-induced calcium activity and the resulting redistribution of biomass within the plasmodial sheet.

## Discussion

Our experimental results, together with our mathematical models, identify a unifying mechanochemical mechanism that links intracellular signalling to organism-scale behaviour in *Physarum poly-cephalum*. In this mechanism, environmentally modulated intracellular calcium oscillations, coupled through diffusion and linked to contractile mechanics, encode and propagate local information across the cell. Experimentally, we show that a local reduction in cortical actin is sufficient to induce mechanical asymmetry and drive directed growth, even in the absence of an external attractant. This suggests that directional behaviour in slime mould can arise from local modulation of cortical mechanics. Building on previous experimental evidence linking calcium concentration to actin depolymerization and contractile relaxation, we propose that environmental cues act through intracellular calcium dynamics to generate these cortical asymmetries. When incorporated into a mechanochemical model, local modulation of calcium oscillations propagate through diffusive coupling and coordinates mechanical activity across the organism, generating a wide range of complex organism-scale behaviours found in *Physarum*: directed migration, network formation, and exploration–exploitation trade-offs.

Recent explanations of long range coordination in *Physarum* often frame the problem of coordination in terms of signalling molecule transport, asking which transport mechanisms can account for the speed of intracellular reorganizations [23, 53]. This perspective implicitly treats coordination as a transport problem, in which signals must traverse the organism via cytoplasmic flow to coordinate behaviour. However, coordination in *Physarum* does not need to rely on long-range transport. By viewing *Physarum* as a spatially extended system of locally coupled biochemical oscillators, coordination instead arises through the propagation of spatiotemporal patterns. In such systems, information travels as waves or synchronized oscillations, moving with a speed set by both the diffusion rate and local reaction kinetics, rather than by long-range molecular signalling transport [54, 55]. In our model, intracellular calcium oscillations provide a biological implementation of such a mechanism, regulating contractility and generating spatio-temporal patterns of mechanical activity across the organism. Thus, we show that cytoplasmic streaming may not be the primary mechanism for coordinating behaviour, but rather a mechanical consequence of oscillatory contraction dynamics that already organize the system at the organism scale. Instead, streaming may mainly serve roles in mass transport, nutrient redistribution, and morphological remodelling, while coordination itself arises from diffusion-coupled biochemical oscillations.

In a sense, our results provide a concrete mechanistic realization of the hypothesis put forward by Boussard et al. who suggested that self-sustained oscillations may sustain learning in slime moulds [4]. This is also consistent with the broader idea that intracellular oscillations are a fundamental function for information processing and distribution in non-neural organisms [56]. From an evolutionary perspective, genomic analyses indicate that *Physarum polycephalum* retains features consistent with an ancestral eukaryotic signalling repertoire, including pathways shared with both Amoebozoa and Opisthokonta [57]. Moreover, calcium is an ancient, evolutionary conserved intracellular signalling molecule across eukaryotes [19, 18, 55]. Thus, the mechanism we identify in our work might represent one possible implementation of a more general mechanism in which oscillatory calcium signalling is coupled to physical processes to coordinate behaviour in living systems. As such, our results can be interpreted in light of hypotheses on the early evolution of nervous systems, which propose that coordination initially relied on diffusion-mediated signalling, often described as volume transmission, before the emergence of synaptic wiring [58, 59]. In this view, spatially extended chemical fields provide a substrate for information processing in the absence of specialised neuronal structures. Our results are also consistent with the perspective that early nervous systems did not primarily evolve as input-output devices, but rather as a way to coordinate contractile activity across organisms, to enable coherent movement [60].

Although our framework captures a broad range of behaviours in *Physarum* within a single mechanochemical model, the behavioural repertoire of slime moulds is even richer. Experiments have demonstrated additional phenomena such as habituation to repeated stimuli [61], anticipation of periodic environmental changes [62], and maze solving [63]. *Physarum*’s ability solve mazes rely on the same processes of pathway formation and pruning as network optimization, but in geometrically constrained environments. As such, maze-solving could be demonstrated with our model implemented on geometrically constrained domains. Experiments show that *Physarum* can anticipate periodic environmental stress, such as cold and dry conditions [62]. This behaviour has been described using phenomenological models of coupled oscillators [62]. If such stressors modulate intracellular calcium dynamics, the oscillatory system in our model could provide a mechanistic basis for this behaviour, with entrainment to periodic inputs giving rise to anticipatory dynamics. Habituation, on the other hand, likely involves adaptive changes in the organism’s signalling pathways [61], which are not captured in our current framework. Capturing this behaviour would require both a better understanding of its biological basis and an explicit, history-dependent representation of how repeated stimulation modifies the coupling between external inputs and intracellular dynamics. While our current model focuses on nutrient cues, represented through the parameter *Q*, corresponding to IP_3_-mediated calcium release, *Physarum* responds to several environmental signals, including light [64] and chemical attractants and repellents [65, 50]. These signals are also known to modulate intracellular calcium dynamics, but may act on distinct components of the calcium oscillatory cycle, for example by altering calcium reuptake, channel inactivation, or baseline calcium levels. Extending the framework to account for such stimulus-specific modulation is thus an important direction for future work. Another natural step is extend our simulations to them to larger domains, which are currently limited by computational costs. Our model also generates experimentally testable predictions. In particular, it suggests that local weakening of actin polymerization through Cytochalasin-D may redistribute biomass without inducing travelling waves, whereas modulation of calcium dynamics through external nutrient stimuli may generate both directed wave propagation and mass redistribution. Testing this distinction experimentally could help assess the relative roles of mechanical and biochemical mechanisms in directed growth and biomass redistribution.

In conclusion, we proposed here a unifying mechanism that bridges the gap between molecular signalling and organism-scale behaviour in *Physarum polycephalum*. Because calcium signalling is both ancient and deeply conserved across eukaryotes, the mechanochemical principle identified here is likely to extend well beyond *Physarum* to explain how complex, coordinated behaviour can emerge in living systems before the evolution of dedicated neural structures. More broadly, our results invite a reframing of biological information processing: rather than being unique to nervous systems, adaptive coordination may be a more general property of oscillatory, mechanically coupled biochemical networks — a fundamental feature of living systems that evolution has repeatedly built upon.

## Materials and Methods

### Experimental Protocol

Tubule-shaped cells of *Physarum* were obtained by placing agar-only and 5% oat-agar blocks ∼ 5cm apart above a water pool. After 18-24 hours, ∼ 2.5 cm tubules were excised from the resulting network and placed in contact with either oat flakes or 1 % non-nutrient agar blocks soaked in 20 *µ*M Cytochalasin-D (CyD) or water. Tubules were exposed to one of four conditions: no treatment (control), asymmetric food (oat flake on one end, water-agar on the other), symmetric food (oat flakes on both ends), or asymmetric CyD (CyD-agar on one end, water-agar on the other). Migratory behaviour was recorded via time-lapse photography (1 frame/min, 12 h) under a 610 nm longpass filter. Final choice was scored by visual inspection as left, right, both, or no choice; “both” and “no choice” were pooled as “undecided.” Behavioural experiments were conducted only for asymmetric conditions. A one-sample proportionality test with Yates’ continuity correction was used to assess directional preference, with undecided trials split equally between the two sides.

For actin imaging, tubules were fixed after 1 hour of treatment exposure, permeabilized, and stained with rhodamine-phalloidin. Confocal Z-stacks (40×) were acquired at four equidistant positions (P1–P4) along each tubule. Stacks were combined per replicate for normalization, then separated and projected as maximum intensity projections (MIPs); mean fluorescence intensity within hand-drawn ROIs was measured in ImageJ. Replicates displaying punctated rather than filamentous actin were excluded. A linear mixed model (LMM) with position and treatment as fixed effects and tubule replicate as a random effect was fit to the normalized intensities; pairwise comparisons used estimated marginal means (EMMs).

### Tubule model

The one-dimensional tubule model (Eqs. (1)–(4)) describes the coupling between intracellular calcium dynamics, active cortical pressure, and tubule radius deformation. Full derivations are provided in the Supplementary Section SI2. The calcium-dependent modulation of contractility is described by a sigmoidal function

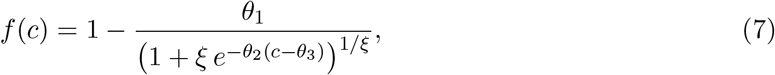

which captures threshold-like actin depolymerization. Parameter values are given in Supplementary Table SI1.

### Phase field model

The phase-field model extends the tubule framework to two spatial dimensions by representing the plasmodium with a phase field *ϕ*(*x, y, t*) and a thickness field *h*(*x, y, t*). The evolution equations for *ϕ* and *h* are given by Eqs. (5) and (6), respectively, in the Results section. In the phase field equation, the double-well potential *G*(*ϕ*) is given by:

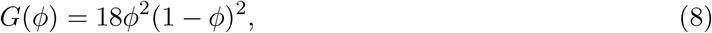

following prior phase-field models of cell movement [42, 41]. This potential drives the phase field toward either 0 or 1, corresponding to outside or inside the cell. The calcium dynamics in the phase field model is given by

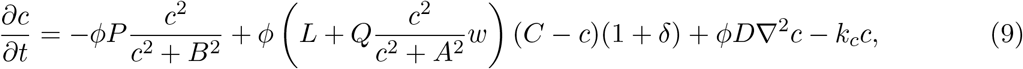

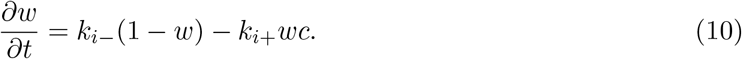

The factors of *ϕ* restrict calcium dynamics to the cell interior, while the term −*k*_*c*_*c* ensures that residual calcium outside the cell decays after the cell has left that region. The gating variable *w* is evolved on the full domain for computational convenience, but only affects the intracellular dynamics. Full derivations are provided in the Supplementary Section SI3.

Numerical simulations were performed using finite-difference methods implemented in Julia. Further details of the numerical implementation and parameter values are provided in the Supplementary Section SI4 and the Supplementary Table SI1, respectively. The code used to generate the results is available at https://github.com/linneagyllingberg/physarum.

## References

[1] František Baluška, Stefano Mancuso, and Dieter Volkmann. Communication in plants. Neuronal aspect of plant life., Spriger Berlin Heidelberg, New York, 2006.

[2] Pamela Lyon. The cognitive cell: bacterial behavior reconsidered. Frontiers in microbiology, 6:264, 2015.

[3] František Baluška and Michael Levin. On having no head: cognition throughout biological systems. Frontiers in psychology, 7:902, 2016.

[4] Auréle Boussard, Adrian Fessel, Christina Oettmeier, Léa Briard, Hans-Günther Döbereiner, and Audrey Dussutour. Adaptive behaviour and learning in slime moulds: the role of oscillations. Philosophical Transactions of the Royal Society B, 376(1820):20190757, 2021.

[5] Marcelo Behar and Alexander Hoffmann. Understanding the temporal codes of intra-cellular signals. Current opinion in genetics & development, 20(6):684–693, 2010.

[6] Jeremy E Purvis and Galit Lahav. Encoding and decoding cellular information through signaling dynamics. Cell, 152(5):945–956, 2013.

[7] Akihiro Isomura and Ryoichiro Kageyama. Ultradian oscillations and pulses: coordinating cellular responses and cell fate decisions. Development, 141(19):3627–3636, 2014.

[8] Randall L Kincaid and Tag E Mansour. Measurement of chemotaxis in the slime mold physarum polycephalum. Experimental cell research, 116(2):365–375, 1978.

[9] Atsushi Tero, Seiji Takagi, Tetsu Saigusa, Kentaro Ito, Dan P Bebber, Mark D Fricker, Kenji Yumiki, Ryo Kobayashi, and Toshiyuki Nakagaki. Rules for biologically inspired adaptive network design. Science, 327(5964):439–442, 2010.

[10] Chris R Reid, Hannelore MacDonald, Richard P Mann, James AR Marshall, Tanya Latty, and Simon Garnier. Decision-making without a brain: how an amoeboid organism solves the two-armed bandit. Journal of The Royal Society Interface, 13(119):20160030, 2016.

[11] Christina Oettmeier, Klaudia Brix, and Hans-Günther Döbereiner. Physarum polycephalum, a new take on a classic model system. Journal of Physics D: Applied Physics, 50(41):413001, 2017.

[12] Karen Alim, Gabriel Amselem, François Peaudecerf, Michael P Brenner, and Anne Pringle. Random network peristalsis in physarum polycephalum organizes fluid flows across an individual. Proceedings of the National Academy of Sciences, 110(33):13306–13311, 2013.

[13] EB Ridgway and AC Durham. Oscillations of calcium ion concentrations in physarum polycephalum. The Journal of cell biology, 69(1):223–226, 1976.

[14] Y Yoshimoto, F Matsumura, and N Kamiya. Simultaneous oscillations of ca2+ efflux and tension generation in the permealized plasmodial strand of physarum. Cell motility, 1(4):433– 443, 1981.

[15] Shinji Yoshiyama, Mitsuo Ishigami, Akio Nakamura, and Kazuhiro Kohama. Calcium wave for cytoplasmic streaming of physarum polycephalum. Cell biology international, 34(1):35–40, 2010.

[16] Y Yoshimoto and N Kamiya. Ca2+ oscillation in the homogenate of physarum plasmodium. Protoplasma, 110(1):63–65, 1982.

[17] David Vogel, Stamatios C Nicolis, Alfonso Perez-Escudero, Vidyanand Nanjundiah, David JT Sumpter, and Audrey Dussutour. Phenotypic variability in unicellular organisms: from calcium signalling to social behaviour. Proceedings of the Royal Society B: Biological Sciences, 282(1819), 2015.

[18] Michael J Berridge, Peter Lipp, and Martin D Bootman. The versatility and universality of calcium signalling. Nature reviews Molecular cell biology, 1(1):11–21, 2000.

[19] R Maynard Case, David Eisner, Alison Gurney, Owen Jones, Shmuel Muallem, and Alexei Verkhratsky. Evolution of calcium homeostasis: from birth of the first cell to an omnipresent signalling system. Cell calcium, 42(4-5):345–350, 2007.

[20] Ernesto Carafoli and Joachim Krebs. Why calcium? how calcium became the best communicator. Journal of Biological Chemistry, 291(40):20849–20857, 2016.

[21] VA Teplov, Yu M Romanovsky, and OA Latushkin. A continuum model of contraction waves and protoplasm streaming in strands of physarum plasmodium. Biosystems, 24(4):269–289, 1991.

[22] Ryo Kobayashi, Atsushi Tero, and Toshiyuki Nakagaki. Mathematical model for rhythmic protoplasmic movement in the true slime mold. Journal of Mathematical Biology, 53(2):273– 286, 2006.

[23] Karen Alim, Natalie Andrew, Anne Pringle, and Michael P Brenner. Mechanism of signal propagation in physarum polycephalum. Proceedings of the National Academy of Sciences, 114(20):5136–5141, 2017.

[24] Atsushi Tero, Ryo Kobayashi, and Toshiyuki Nakagaki. Physarum solver: A biologically inspired method of road-network navigation. Physica A: Statistical Mechanics and its Applications, 363(1):115–119, 2006.

[25] Atsushi Tero, Ryo Kobayashi, and Toshiyuki Nakagaki. A mathematical model for adaptive transport network in path finding by true slime mold. Journal of theoretical biology, 244(4):553– 564, 2007.

[26] Linnéa Gyllingberg, Yu Tian, and David JT Sumpter. A minimal model of cognition based on oscillatory and current-based reinforcement processes. Journal of the Royal Society Interface, 22(222):rsif–2024, 2025.

[27] Farshad Ghanbari, Joe Sgarrella, and Christian Peco. Emergent dynamics in slime mold networks. Journal of the Mechanics and Physics of Solids, 179:105387, 2023.

[28] Takayuki Hasegawa, Sho Takahashi, Hiroshi Hayashi, and Sadashi Hatano. Fragmin: a calcium ion sensitive regulatory factor on the formation of actin filaments. Biochemistry, 19(12):2677– 2683, 1980.

[29] Hiroaki Sugino and Fumio Matsumura. Fragmin induces tension reduction of actomyosin threads in the presence of micromolar levels of ca2+. The Journal of cell biology, 96(1):199– 203, 1983.

[30] Kazuhiro Kohama. Calcium inhibition as an intracellular signal for actin–myosin interaction. Proceedings of the Japan Academy, Series B, 92(10):478–498, 2016.

[31] Tsuyoshi Okagaki, Sugie Higashi-Fujime, and Kazuhiro Kohama. Ca2+ activates actinfilament sliding on scallop myosin but inhibits that on physarum myosin. The Journal of Biochemistry, 106(6):955–957, 1989.

[32] Kazuhiro Kohama and Setsuro Ebashi. Inhibitory ca2+-regulation of the physarum actomyosin system. In The molecular biology of Physarum polycephalum, pages 175–190. Springer, 1986.

[33] Michael J Berridge. The inositol trisphosphate/calcium signaling pathway in health and disease. Physiological reviews, 96(4):1261–1296, 2016.

[34] Sophie Marbach, Noah Ziethen, and Karen Alim. Vascular adaptation model from force balance: Physarum polycephalum as a case study. New Journal of Physics, 25(12):123052, 2023.

[35] J Poledna. Model of intracellular calcium oscillations activated by inositol trisphosphate. General physiology and biophysics, 12:381–381, 1993.

[36] Bjoern Kscheschinski, Mirna Kramar, and Karen Alim. Calcium regulates cortex contraction in physarum polycephalum. Physical Biology, 21(1):016001, 2023.

[37] Jean-Paul Rieu, Hélene Delanoë-Ayari, Seiji Takagi, Yoshimi Tanaka, and Toshiyuki Nakagaki. Periodic traction in migrating large amoeba of physarum polycephalum. Journal of The Royal Society Interface, 12(106):20150099, 2015.

[38] Yoshihiro Miyake, Hideki Tada, Masafumi Yano, and Hiroshi Shimizu. Relationship between intracellular period modulation and external environment change in physarum plasmodium. Cell structure and function, 19(6):363–370, 1994.

[39] Makiko Nonomura. Study on multicellular systems using a phase field model. PloS one, 7(4):e33501, 2012.

[40] Adrian Moure and Hector Gomez. Phase-field modeling of individual and collective cell migration. Archives of Computational Methods in Engineering, 28(2):311–344, 2021.

[41] Sergio Alonso, Maike Stange, and Carsten Beta. Modeling random crawling, membrane deformation and intracellular polarity of motile amoeboid cells. PloS one, 13(8):e0201977, 2018.

[42] Danying Shao, Wouter-Jan Rappel, and Herbert Levine. Computational model for cell morphodynamics. Physical review letters, 105(10):108104, 2010.

[43] Danying Shao, Herbert Levine, and Wouter-Jan Rappel. Coupling actin flow, adhesion, and morphology in a computational cell motility model. Proceedings of the National Academy of Sciences, 109(18):6851–6856, 2012.

[44] EN Brewer, Susumu Kuraishi, Joseph C Garver, and FM Strong. Mass culture of a slime mold, physarum polycephalum. Applied Microbiology, 12(2):161–164, 1964.

[45] Werner Baumgarten and Marcus JB Hauser. Dynamics of frontal extension of an amoeboid cell. Europhysics Letters, 108(5):50010, 2014.

[46] Tetsuo Ueda, Yoshihito Mori, and Yonosuke Kobatake. Patterns in the distribution of intracellular atp concentration in relation to coordination of amoeboid cell behavior in physarum polycephalum. Experimental cell research, 169(1):191–201, 1987.

[47] DR Coman. Additional observations on positive and negative chemotaxis: experiments with a myxomycete. Arch. Pathol, 29:220–228, 1940.

[48] Randall L Kincaid and Tag E Mansour. Chemotaxis toward carbohydrates and amino acids in physarum polycephalum. Experimental Cell Research, 116(2):377–385, 1978.

[49] K Natsume, Y Miyake, M Yano, and H Shimizu. Development of spatio-temporal pattern of ca2+ on the chemotactic behaviour of physarum plasmodium. Protoplasma, 166(1):55–60, 1992.

[50] Kenji Matsumoto, Seiji Takagi, and Toshiyuki Nakagaki. Locomotive mechanism of physarum plasmodia based on spatiotemporal analysis of protoplasmic streaming. Biophysical journal, 94(7):2492–2504, 2008.

[51] Audrey Dussutour and Chloé Arson. Flow-network adaptation and behavior in slime molds. Fungal Ecology, 68:101325, 2024.

[52] Subash Kusum Ray. Brainless. but Smart: Investigating Cognitive-Like Behaviors in the Acelular Slime Mold Physarum Polycephalum. PhD thesis, New Jersey Institute of Technology, 2022.

[53] Johnny Tong, Kaspar Wachinger, Fabian K Henn, Nico Schramma, Siyu Chen, and Karen Alim. Coexistence of trapped and flow-transported nuclei enables fast pigeon post communication across multinucleated cell. Proceedings of the National Academy of Sciences, 122(50):e2411101122, 2025.

[54] James Dickson Murray and James Dickson Murray. Mathematical biology: II: spatial models and biomedical applications, volume 18. Springer, 2003.

[55] Martin Falcke. Reading the patterns in living cells—the physics of ca2+ signaling. Advances in physics, 53(3):255–440, 2004.

[56] Pamela Lyon, Fred Keijzer, Detlev Arendt, and Michael Levin. Reframing cognition: getting down to biological basics. Philosophical Transactions of the Royal Society B, 376(1820):20190750, 2021.

[57] Pauline Schaap, Israel Barrantes, Pat Minx, Narie Sasaki, Roger W Anderson, Marianne Bénard, Kyle K Biggar, Nicolas E Buchler, Ralf Bundschuh, Xiao Chen, et al. The physarum polycephalum genome reveals extensive use of prokaryotic two-component and metazoan-type tyrosine kinase signaling. Genome Biology and Evolution, 8(1):109–125, 2016.

[58] Gáspár Jékely. The chemical brain hypothesis for the origin of nervous systems. Philosophical Transactions of the Royal Society B: Biological Sciences, 376(1821), 2021.

[59] Leonid L Moroz. On the independent origins of complex brains and neurons. Brain Behavior and Evolution, 74(3):177–190, 2009.

[60] Fred Keijzer, Marc Van Duijn, and Pamela Lyon. What nervous systems do: early evolution, input–output, and the skin brain thesis. Adaptive Behavior, 21(2):67–85, 2013.

[61] Romain P Boisseau, David Vogel, and Audrey Dussutour. Habituation in non-neural organisms: evidence from slime moulds. Proceedings of the Royal Society B: Biological Sciences, 283(1829):20160446, 2016.

[62] Tetsu Saigusa, Atsushi Tero, Toshiyuki Nakagaki, and Yoshiki Kuramoto. Amoebae anticipate periodic events. Physical review letters, 100(1):018101, 2008.

[63] Toshiyuki Nakagaki, Hiroyasu Yamada, and Ágota Tóth. Maze-solving by an amoeboid organism. Nature, 407(6803):470–470, 2000.

[64] Donat Häder. Role of calcium in phototaxis of physarum polycephalum. Plant and cell physiology, 26(7):1411–1417, 1985.

[65] T Ueda, M Muratsugu, K Kurihara, and Y Kobatake. Chemotaxis in physarum polycephalum: Effects of chemicals on isometric tension of the plasmodial strand in relation to chemotactic movement. Experimental cell research, 100(2):337–344, 1976.

