## Supplementary Information for "Self-organizing physical and biochemical interactions explain diverse behaviours in *Physarum polycephalum*"

Linnéa Gyllingberg[1,*,†], Abid Haque[2,3,†], Subash K. Ray[2,3],

Gregory Weber[3,4], Jason M. Graham[5], Simon Garnier[2,3]

[1]Department of Mathematics, University of California, Los Angeles, CA, USA
[2]Federated Department of Biological Sciences, New Jersey Institute of Technology, Newark, NJ, USA
[3]Federated Department of Biological Sciences, Rutgers University-Newark, Newark, NJ, USA
[4]Department of Biology, University of Indianapolis, Indianapolis, IN, USA
[5]Department of Mathematics, University of Scranton, Scranton, PA, USA
[†]L.G. and A.H. contributed equally to this work.

#### Introduction

This appendix is organised as follows. Section SI1 describes the experimental setup and data analysis. Section SI2 derives the one-dimensional tubule model, and Section SI3 extends the framework to the phase-field model. Section SI4 summarizes the numerical implementation and parameter values. Finally, Section SI5 defines the quantitative metrics used to analyse the simulations and presents the corresponding supplementary figures.

#### SI1 Experiments

##### SI1.1 Experimental setup

For both the behavioural experiments and actin imaging, we used tubule-shaped cells of *Physarum* which were obtained using the following protocol. We placed two agar blocks – one containing only agar, and the other containing 5% oat-agar (or food agar) – on a support above a pool of water. The distance between the two blocks was approximately 5 cm. A biomass of *Physarum* weighing approximately 0.5 g was placed on top of the agar-only block and allowed to grow tubular extensions on the surface of the water. After 18-24 hours, a tubule network had formed between the agar-only and food-agar blocks. Tubules of approximately 2.5 cm in length were excised from the network, straightened, and placed in contact with oat flakes (food experiments), or agar blocks soaked with Cytochalasin-D or water in the final experimental setup (Fig. SI1). The agar blocks were prepared using 1 % non-nutrient agar, cut into blocks of approximate size $2\,\mathrm{mm}^2$. These blocks were subsequently stored in tubes either containing water, or a 20 $\mu$M

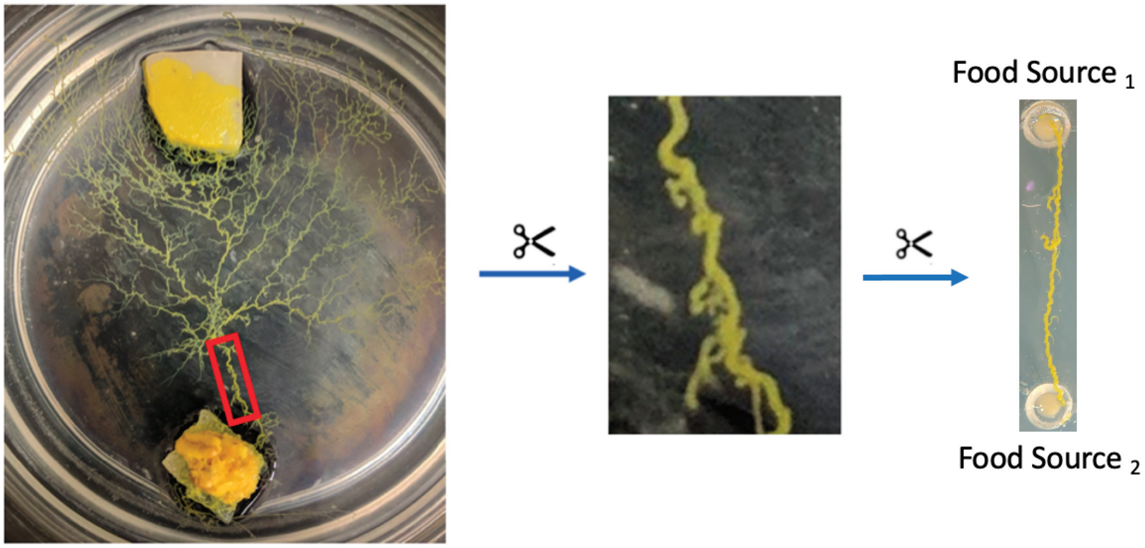


Fig. SI1: Tubules, approximately 2 cm long, excised from a larger slime mould network (red rectangle) were used for studying cytoskeletal structure and dynamics. In the actual experiment, Food $\text{source}_1$ and Food $\text{source}_2$ refer to treatment substances between which the slime mould makes a choice (food/$H_2O$/Cytochalasin D).

Cytochalasin-D solution. To allow for the uniform soaking of the agar blocks, we stored the agar blocks in the respective solutions over 24 hours in a $-4°$C incubator.

In our final experimental setup, we used a Petri plate with a diameter of 6 cm and height of 1.3 cm. A 1% w/v non-nutrient agar substrate was used to provide a gel-like and moist base. The *Physarum* tubule was exposed to four choice treatments. In the first treatment, a tubule was placed on the experimental arena, with zero exposure to food or any other reagents (referred to as the Control condition). In the second treatment, one oat flake soaked with room temperature water was placed at one end of the tubule (referred to as the "asymmetric food choice" condition). In the third treatment, two oat flakes soaked with room temperature water were placed, one at the left end and the other at right end of the tubule (referred to as the "symmetric food choice"). Finally, in the fourth treatment, we placed the tubule in contact with an agar block soaked with 20 $\mu$M Cytochalasin-D obtained from ThermoFisher Scientific® on one end, and an agar block soaked with water on the other end (referred to as the "CyD choice condition"). We recorded the migratory behavior of the *Physarum* tubule using high-resolution time-lapse photography. A Panasonic® Lumix GH4 camera was used in combination with an Olympus® M.Zuiko Digital ED 60mm macro lens to capture one image every minute for a total of 12 hours. To obtain high-resolution images of the tubule, we only recorded the portion of the experimental setup where the tubules were situated, and the camera was zoomed in as closely as possible. The experimental setup was illuminated from below using an LED panel from www.superbrightleds.com®, which allowed for the recording of high-definition tubule edges. To prevent *Physarum* from being sensitive to and avoiding UV and short-wavelength visible light, a 610 nm longpass filter from Newport Corporation® was placed between the experimental setup and the LED panel. Previous studies have demonstrated that *Physarum* is not sensitive to wavelengths of light that pass through this filter [1].

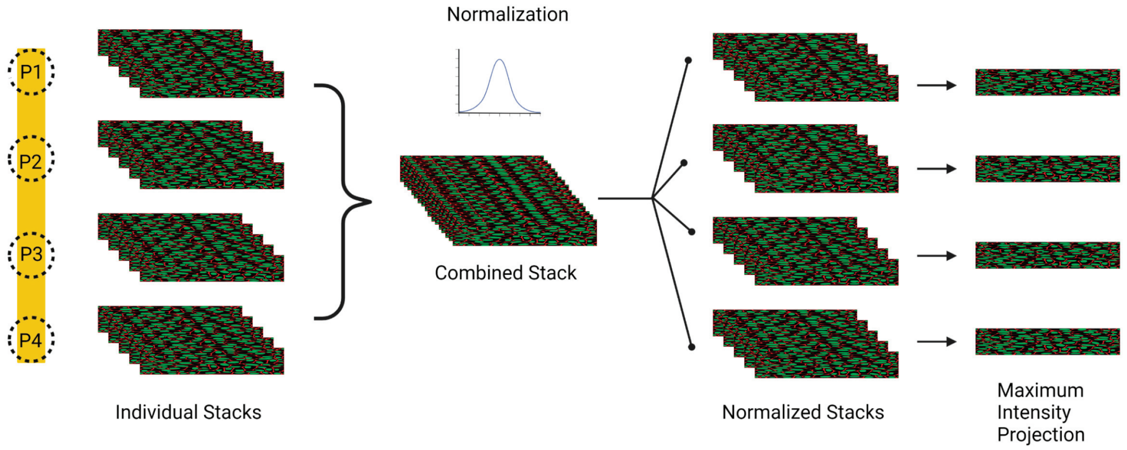


Fig. SI2: Standardized procedure for comparing actin fluorescence intensity between four equidistant locations on a tubule. First, image stacks were collected from each location. These stacks were then combined into one single stack per replicate, and normalized. Then the stacks were separated, and a maximum intensity projection (MIP) was obtained for each individual location. The mean fluorescence intensity from each MIP was used for further statistical comparison.

The use of opaque oat flakes and transparent agar blocks which refracted the background light made it difficult for a computer vision software to measure the growth of the slime mould tubule on the treatment substance. Therefore, we conducted a visual inspection of the experimental setup at the end of the experiment to determine the final food choice of the *Physarum* tubule. If a discernible amount of biomass aggregation was observed exclusively on the left or right treatment substance (oat flake, water or cytochalasin-D), the final choice was recorded as "left" or "right", respectively. If the relative growth of P. polycephalum biomass on the treatment substances was visually indistinguishable, the final outcome of the experiment was recorded as "both". If the tubule retracted or broke contact with both of the offered choices, the experiment was recorded as "no choice". Both the "both" and "no choice" results were interpreted together as "undecided", as the tubule fails to show a preference for either of the food sources. The behavioural choice experiments were only performed for the asymmetric choice conditions, since symmetric choice conditions have already been shown to elicit an insignificant preference for either of the treatment sources [2].

To investigate the role of the actin cytoskeleton in the decision-making process of *Physarum* in choosing a treatment substance, we conducted the following experimental protocol. We started the experiment by putting an initially unpolarized *Physarum* tubule in physical contact with no oat flakes for the control condition, one oat flake on one side for the asymmetric food choice condition, one oat flake soaked with room temperature water on each side for the symmetric food choice condition, and one agar block soaked with Cytochalasin-D and the other soaked with water for the CyD choice condition. All the cells were exposed to their respective conditions for a duration of 1 hour in a dark enclosure at room temperature. This allowed the cell to interact with the treatment substance and undergo the decision-making process.

#### SI1.2 Metrics

For the behavioural choice experiments, we used a one-sample proportionality test with Yates' continuity correction [3] to determine whether the tubules had a statistically significant preference for either of the treatments offered on each side. Under the null hypothesis (i.e., no preference for either of the two treatment substances), we expect the theoretical proportions of choosing either of the two food sources to be equal. However, calculating the theoretical proportion of undecided cases (where the tubules chose either both or none of the agar blocks) requires knowledge of the exact decision-making mechanism, which is currently unknown. Nonetheless, the undecided cases are an important aspect of the data because they indicate when *Physarum* was unable to display a preference for either of the treatment substances. Therefore, half of the undecided experimental trials were counted towards the left treatment substance, and the other half were counted towards the right treatment substance, to ensure that the lack of choice in one or more trials was reflected in the final outcome of the analysis. This approach ensured that the undecided cases were not discarded from the analysis, and that the final outcome accurately represented the behaviour of *Physarum polycephalum* in response to the treatment substances presented to it.

To quantify the abundance of actin at different locations on a *Physarum* tubule, we performed the following protocol. First, four image stacks of the tubule were captured from four equidistant locations using a confocal microscope. These locations were chosen based on their proximity to the food sources, as well as their distance from each other. These image stacks were then combined into a single stack for normalization. We normalized the fluorescence intensity of the actin signal in each experimental replicate to account for variations in the staining and imaging process. The normalized stack was then separated into individual stacks representing the four locations on a tubule, and a maximum intensity projection (MIP) was generated to obtain a two-dimensional image of the actin cortex (Fig. SI2). Next, regions of interest (ROIs) were selected from these MIPs, which excluded the background. The mean intensity of fluorescence within each ROI was then measured using ImageJ software. This analysis provided a summary statistic of the abundance of actin at each location on each tubule replicate. For the next steps of analysis, we normalized the mean intensity values of the replicates within each treatment condition with respect to the maximum mean fluorescence intensity observed for that treatment condition. We performed this normalization to eliminate differences in the baseline fluorescence intensity observed in each treatment, so that the analysis can be focused on the comparison of the spatial patterns of mean actin fluorescence intensity between the different treatments. It must be noted here that some replicates displayed a highly punctated organization of actin, instead of a fiber-like organization. The punctates typically had high fluorescence values, which can occur because of an aggregation of actin, or because of an aggregation of fluorescent molecules. The high fluorescence intensity values of such replicates interfere with the normalization procedure, as it suppresses the fluorescence intensity values of non-punctated replicates. Hence, these replicates were excluded from the normalization procedure and not analyzed further.

To study the relationship between the abundance of actin and the distance from the treatment substance, we fit a Linear Mixed Model (LMM) to our data, with the locations (P1, P2, P3 and P4) and treatment (control, symmetric food choice, asymmetric food choice and asymmetric cytochalasin D choice) as fixed effects, and the tubule replicate as a random effect (Fig. SI3). We treated the tubule replicate as a random effect because measurements of actin fluorescence intensity on the same tubule cannot be considered as independent from each other. This allowed us to account for the variability within and between tubules, and to estimate the dependence of

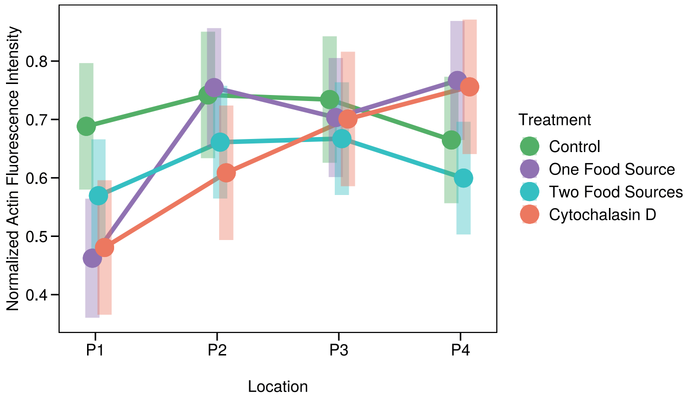


Fig. SI3: Estimated marginal means of normalized actin fluorescence intensity across locations along the tubule (P1–P4) for each treatment condition. Points show model-estimated means from a linear mixed model with location, treatment, and tubule replicate as a random intercept. Error bars show confidence intervals. In the one-food-source and cytochalasin-D treatments, the treatment substance was placed at P1; in the two-food-source treatment, food sources were placed at both P1 and P4.

mean fluorescence intensity on the spatial distance from a treatment substance. Specifically, we used the LMM to assess whether the abundance of actin varied significantly at different distances from the food sources, and to quantify the magnitude and direction of this variation. To account for the effects of the interacting independent variables, we used estimated marginal means. Estimated marginal means are a useful tool for interpreting the effects of multiple independent variables on a response variable, as they provide a summary of the response variable adjusted for the effects of all other variables in the model. In our case, we used estimated marginal means to determine the relationship between the abundance of actin and the distance from the food sources, while accounting for the effects of the treatment substance. By using estimated marginal means, we were able to more accurately assess the effects of the independent variables on the response variable, and to draw more reliable conclusions about the mechanisms that govern the decision-making process of the slime mould cell in response to food sources and Cytochalasin-D.

### SI2 Derivation of the tubule model

We model the cell as a closed hydraulic and chemical system interacting with its environment, building on existing models of slime mould mechanics [4] and $IP_3$-activated calcium dynamics [5]. The aim is to connect short-timescale intracellular oscillations to longer-timescale biomass redistribution, symmetry breaking, and morphological change in *Physarum polycephalum*. The central idea is that local biochemical signalling, cortical force generation, and mechanics are coupled across scales to generate global adaptive behaviour.

#### SI2.1 Radius equations

We start by representing an isolated *Physarum* tubule as a one-dimensional continuous tube of finite length $L$ and variable radius $r(x, t)$, where $x \in [0, L]$, enclosing incompressible cytoplasm. The tube wall is assumed to be viscoelastic and capable of active contraction. Following Teplov et al. [4], we assume that adhesion to the substrate suppresses longitudinal deformation, so that contractility primarily induces local radial changes, and do not alter the length of the tubule. Under these assumptions, the radius dynamics are governed by the passive wall mechanics, the viscous resistance of the endoplasm, and the active pressure generated by the cortex:

$$\frac{\partial r}{\partial t} = \frac{R\eta d}{16\mu}\frac{\partial}{\partial t}\nabla^2 r + \frac{RE_1 d}{16\mu}\nabla^2 r + \frac{R^3}{16\mu}\nabla^2 P^{(A)}. \tag{SI1}$$

Here, $R$ is the mean tubule radius, $d$ the cortical wall thickness, $\mu$ the dynamic viscosity of the endoplasm, $\eta$ the viscous coefficient of the cortical wall, and $E_1$ its elastic stiffness. The first two terms describe the passive viscoelastic and elastic response of the wall, whereas the third term captures active deformation due to gradients in cortical active pressure.

#### SI2.2 Pressure equation

While the continuum model introduced by Teplov et al. [4] captures the mechanical response of a deformable tubule, the active pressure term in that formulation is derived by implicitly assuming a muscle-like mechanism of force generation. In this formulation, the density of actin filaments is treated as approximately constant and contraction arises primarily from myosin-driven sliding of parallel actin filaments.

However, experimental studies of *Physarum* show that cortical force generation can be understood as a combination of the polymerization state of actin, and the reversible cyclic cross-linking of actin filaments by myosin. In particular, periodic contraction–relaxation cycles have been associated with cyclic polymerization and depolymerization of cortical actin filaments and their reversible cross-linking by myosin motors [6, 7]. These observations suggest that active stress should depend on the local polymerization state of the actin network, that is, whether actin is polymerized or depolymerized, rather than on a fixed actin network with variable motor activity.

Thus, we couple actin polymerization dynamics to the active mechanical stress generated in the slime mould tubule. Because actin polymerization is reversible, and filament growth occurs at uncapped barbed ends that can be blocked by capping proteins, the actin network can be described, at a coarse-grained level, by three mechanically relevant states: globular actin monomers ($G$), uncapped filamentous actin ($F_u$) with free barbed ends that can incorporate additional monomers, and capped filamentous actin ($F_b$) whose barbed ends are blocked and can no longer accept new monomers.

The capped filaments form the mechanically stable actin network at the cortex and therefore contribute directly to the generation of contractile stress. We can thus represent the cortical actin dynamics by the following minimal mechano-chemical cycle:

$$G + F_u \underset{k_2}{\overset{k_1'}{\rightleftharpoons}} F_b, \tag{SI2}$$

where $G$ denotes globular actin monomers, $F_u$ uncapped filamentous actin with free barbed ends, and $F_b$ capped filamentous actin. The parameters $k_1'$ and $k_2$ denote the forward and backward rate constants for the conversion of G-actin into force-bearing F-actin, respectively.

Since the actin network is distributed along the tubule, the concentrations of these species depend on both space and time. The local concentration of capped filaments therefore evolves according to the reaction kinetics

$$\frac{\partial F_b}{\partial t} = k_1' G F_u - k_2 F_b. \tag{SI3}$$

To reduce this system, we introduce two conservation relations. First, we assume that the total pool of actin monomers available for polymerization is fixed, so that $G = G_0 - F_b$, where $G_0$ is the total monomer pool. Second, we assume that the number of available uncapped filament ends is limited by a maximal filament concentration $F_0$, so that $F_u = F_0 - F_b$. Substituting these expressions into the kinetic equation gives

$$\frac{\partial F_b}{\partial t} = k_1'(G_0 - F_b)(F_0 - F_b) - k_2 F_b. \tag{SI4}$$

We now assume that the pool of monomeric G-actin is much larger than the concentration of capped filaments under physiological conditions, i.e. $G_0 \gg F_b$. Under this assumption, $G_0 - F_b \approx G_0$, while the number of available uncapped filament ends remains limited by $F_0 - F_b$. Equation (SI4) therefore reduces to

$$\frac{\partial F_b}{\partial t} \approx k_1' G_0 (F_0 - F_b) - k_2 F_b.$$

Defining $k_1 = k_1' G_0$ and $F_b^{(0)} = F_0$, this can be written as

$$\frac{\partial F_b}{\partial t} = k_1(F_b^{(0)} - F_b) - k_2 F_b. \tag{SI5}$$

Here, $k_1$ is an effective rate constant for the restoration of force-bearing cortical actin, and $F_b^{(0)}$ denotes the maximum concentration of bounded F-actin with capped ends.

Furthermore, we assume that the tangential active stress, $\sigma_A(x,t)$, is proportional to the concentration of bounded F-actin with capped ends. Defining $\sigma_0$ as the maximum stress corresponding to complete polymerization of the available actin pool, we write

$$\sigma_A = \sigma_0 \frac{F_b}{F_b^{(0)}}.$$

Equivalently, $F_b = F_b^{(0)} \sigma_A / \sigma_0$, and therefore

$$\frac{\partial F_b}{\partial t} = \frac{F_b^{(0)}}{\sigma_0} \frac{\partial \sigma_A}{\partial t}.$$

Substituting this relation into Equation (SI5) gives

$$\frac{\partial \sigma_A}{\partial t} = k_1(\sigma_0 - \sigma_A) - k_2 \sigma_A, \tag{SI6}$$

where $k_1$ and $k_2$ are effective rate constants describing the restoration and loss of force-bearing cortical actin, respectively.

Experimental studies show that $Ca^{2+}$-sensitive proteins such as fragmin sever actin filaments in *Physarum* [8, 9]. This activity is negligible at low $Ca^{2+}$ concentrations, emerges around $10^{-7}$ M, and saturates at higher concentrations. Since we are primarily interested in the oscillatory dynamics of plasmodial tubules, we approximate the underlying actin-regulatory chemistry by a single rate-controlling step. We assume that $Ca^{2+}$ serves as the rate-controlling substance, and incorporate its role in regulating actin depolymerization by allowing the effective rate constant for restoration of force-bearing cortical actin, $k_1$, to depend on the local $Ca^{2+}$ concentration:

$$k_1(c) = k_1 f(c),$$

where $c$ denotes the $Ca^{2+}$ concentration.

To capture the threshold-like relationship between $Ca^{2+}$ concentration and cortical tension, we choose $f(c)$ to be a monotonically decreasing sigmoidal function,

$$f(c) = 1 - \frac{\theta_1}{\left(1 + \xi\, e^{-\theta_2(c-\theta_3)}\right)^{1/\xi}}. \tag{SI7}$$

With this choice, increasing $Ca^{2+}$ reduces the effective restoration of force-bearing cortical actin, capturing the rapid loss of cortical tension that occurs when $Ca^{2+}$ exceeds a threshold concentration. The active stress equation therefore becomes

$$\frac{\partial \sigma_A}{\partial t} = k_1 f(c)(\sigma_0 - \sigma_A) - k_2 \sigma_A. \tag{SI8}$$

Following Teplov et al., we assume that the thickness of the tubule wall $d$ is much smaller than the tubule radius $R$ ($d \ll R$), so that a thin-wall approximation can be used. In this limit, the active pressure $P^{(A)}$ generated by the cortex is related to the tangential active stress $\sigma_A$ by

$$P^{(A)} = \frac{d}{R}\sigma_A.$$

Now defining the maximum active pressure as $P_0^{(A)} = \frac{d}{R}\sigma_0$, corresponding to complete polymerization of the available actin pool, we obtain the following dynamical equation for the active pressure:

$$\frac{\partial P^{(A)}}{\partial t} = k_1 f(c)(P_0^{(A)} - P^{(A)}) - k_2 P^{(A)}. \tag{SI9}$$

This has the same reduced mathematical form as the active-pressure equation in Teplov et al. [4], but follows from a different interpretation of the force-generating mechanism. Teplov et al. assume that actin filament density remains approximately constant and that active stress arises from myosin binding and sliding on this actin scaffold. Here, following the actin-cycle formulation above, we instead assume that tangential active stress is proportional to the concentration of bounded F-actin with capped ends.

#### SI2.3 Calcium equations

While the model presented by Teplov et al. [4] assumes that calcium release is regulated by mechanical strain, experimental studies suggest that calcium oscillations in *Physarum* can arise autonomously. In particular, oscillations have been observed in slime-mould homogenates, indicating that the calcium dynamics are generated by an intrinsic biochemical oscillator rather than being driven by mechanical deformation [10]. Thus, in our formulation, calcium acts as an upstream regulatory signal that modulates the actin network and contractile stress, while the mechanics does not directly feed back into the calcium dynamics.

Mechanically independent calcium oscillations are a characteristic that *Physarum* shares with a wide range of eukaryotic cells, including plants [11, 12], animals [13, 14], and fungi [15]. As in metazoan cells, calcium in *Physarum* exists both as free cytosolic $Ca^{2+}$ and in internal stores, and external chemical cues are thought to regulate intracellular calcium dynamics through evolutionarily conserved receptor-mediated signalling pathways [16]. In *Physarum*, chemical stimuli such as food are thought to activate G-protein coupled receptor (GPCR)-mediated signalling, leading to the production of inositol 1,4,5-triphosphate ($IP_3$), which triggers calcium release from internal stores into the cytosol [17]. Elevated cytosolic calcium further promotes calcium release through calcium-induced calcium release (CICR), forming a positive feedback loop [18]. In parallel, ATP-dependent pumps transport calcium back into internal stores, providing a negative feedback mechanism that counteracts excessive cytosolic calcium accumulation [19]. The interplay between release and sequestration can therefore generate sustained intracellular calcium oscillations.

To model this, we adopt the phenomenological two-compartment calcium oscillator proposed by Poledna [5], which describes $IP_3$-dependent calcium release, calcium-dependent amplification, and reuptake into internal stores:

$$\frac{\partial c}{\partial t} = -P\,\frac{c^2}{c^2+B^2} + \Big(L + Q(x,t)\,\frac{c^2}{c^2+A^2}\,w\Big)(C-c)(1+\delta) + D\,\nabla^2 c, \qquad \text{(SI10)}$$

$$\frac{\partial w}{\partial t} = k_{i-}(1-w) - k_{i+}\,w\,c, \qquad \text{(SI11)}$$

where $c(x,t)$ denotes the cytosolic calcium concentration and $w(x,t)$ the fraction of calcium release channels available for activation. The calcium dynamics is governed by the exchange of calcium between the cytoplasm and the endoplasmic reticulum, the primary calcium storage. In this set of equations, $P$ denotes the maximum rate of the calcium pump, B sets the calcium concentration at which the pump transitions from low to high activity. $L$ denotes constant leak permeability representing passive calcium release from internal stores, $A$ sets the calcium concentration at which the release channel becomes significantly activated, and $\delta$ is the ratio of effective volumes of calcium storage of the cytoplasm and the reticulum.

In the original formulation, Poledna considered a spatially homogeneous intracellular oscillator. Since our aim is to model calcium dynamics along a spatially extended tubule, and cytosolic calcium is mobile and can spread over short distances through diffusion, we include a diffusion term to account for local spatial coupling of calcium dynamics between neighbouring regions [20], where $D$ is the diffusion constant.

A key parameter in the calcium-release term is $Q(x,t)$, which denotes the maximal permeability of the $Ca^{2+}$-release channels. In *Physarum*, food and other chemical cues are thought to elevate intracellular $Ca^{2+}$ through receptor-mediated signalling [21]. More generally, food-induced stimulation is a well-known trigger of $IP_3$-mediated $Ca^{2+}$ release in diverse cell types [22, 23]. We therefore represent external stimulation in the model as a local increase in $Q$ above its baseline level. In this way, $Q$ serves as the interface between environmental input and intracellular calcium release: larger values of $Q$ increase the potential calcium-release flux through non-inactivated channels, whereas smaller values reduce it. In the one-dimensional tubule simulations, stochastic fluctuations are added around the prescribed spatial profile of $Q(x,t)$ to represent variability in environmental sensing and intracellular signalling.

### SI3 Derivation of the phase-field model

The one-dimensional tubule model captures how intracellular calcium oscillations couple to active pressure and local deformation, but it cannot represent whole-cell migration or large-scale morphological rearrangements. To describe these processes, we extend the framework to two spatial dimensions using a phase-field formulation [24], in which the moving cell boundary is represented implicitly by a continuous field. This allows intracellular biochemical and mechanical dynamics to be solved on a fixed spatial grid [25].

#### SI3.1 Diffuse-interface representation of the plasmodium

We represent the plasmodium by a scalar phase field $\phi(x, y, t) \in [0, 1]$, which describes the local presence of cytoplasm. Regions with $\phi \approx 1$ correspond to the interior of the cell, regions with $\phi \approx 0$ correspond to the exterior, and the cell boundary is represented by a narrow transition region between these states. This diffuse interface provides a well-defined domain for intracellular processes without requiring explicit boundary tracking.

##### SI3.1.1 Phase-field evolution equation

The evolution of the cell boundary is described by

$$\tau \frac{\partial \phi}{\partial t} = f(c)\,\gamma \left( \nabla^2 \phi - \frac{G'(\phi)}{\epsilon^2} \right) - \beta \left( \int \phi\, h\, dx\, dy - V_0 \right) |\nabla \phi| + f(c)\,\alpha |\nabla \phi|. \quad \text{(SI12)}$$

Here, $\tau$ sets the timescale of boundary motion, $\gamma$ is the passive interfacial tension, $\epsilon$ controls the width of the diffuse interface, $\beta$ determines the strength of volume conservation, and $\alpha$ sets the magnitude of active boundary forcing.

This equation is based on three physically motivated contributions. The first term,

$$f(c)\,\gamma \left( \nabla^2 \phi - \frac{G'(\phi)}{\epsilon^2} \right),$$

represents passive interfacial relaxation. We take the double-well potential, $G(\phi)$, following prior phase field formulations for cell movement [26, 27]:

$$G(\phi) = 18\phi^2(1 - \phi)^2, \quad \text{(SI13)}$$

which gives two preferred states, $\phi = 0$ and $\phi = 1$, corresponding to the exterior and interior of the cell. The Laplacian term, $\nabla^2 \phi$, smooths the transition between these two states. Together, these terms produce a cell with a diffuse interface of finite thickness. The parameter $\gamma$ controls the effective interfacial tension. We modulate this term by $f(c)$ to reflect that elevated calcium weakens cortical actin structure and reduces effective cortical tension. The second term,

$$-\beta \left( \int \phi\, h\, dx\, dy - V_0 \right) |\nabla \phi|,$$

enforces approximate conservation of total cell volume. The quantity

$$\int \phi\, h\, dx\, dy$$

represents the total biomass of the plasmodial sheet, and $V_0$ is the reference volume. Deviations from this reference generate a restoring force localized at the boundary through the factor $|\nabla \phi|$. The third term,

$$f(c)\,\alpha |\nabla \phi|,$$

represents active boundary forcing generated by the cortical actomyosin network. Since $|\nabla\phi|$ is zero inside the cell, but non-zero only in the interface region, this term acts locally at the cell boundary. Its modulation by $f(c)$ reflects calcium-dependent weakening of force-bearing cortical actin, so that elevated calcium reduces the effective active boundary forcing. Here, $f(c)$ as defined in the same way as for tubule model, given by Eq. (SI7).

##### SI3.2 Thickness dynamics

Although the model is defined on a two-dimensional domain, redistribution of biomass is captured through a thickness field $h(x, y, t)$, effectively giving a three-dimensional description of the cell:

$$\frac{\partial h}{\partial t} = -\phi f(c) k_{hc} h - \phi k_{hv} \left( \int \phi h \, dx \, dy - V_0 \right) - k_{h-} h. \tag{SI14}$$

The first term represents calcium-modulated contraction, which reduces local thickness. The second term enforces approximate conservation of total biomass. The third term represents passive relaxation of thickness in regions where the plasmodium has retracted.

Having these two distinct variables, the phase field, which defines the cell boundary, and the thickness field, which represents local biomass, allows the model to capture both changes in cell shape and redistribution of mass within the cell.

##### SI3.3 Calcium dynamics in the moving domain

Intracellular calcium dynamics are described by the same oscillator used in the one-dimensional model, extended to two spatial dimensions:

$$\frac{\partial c}{\partial t} = -\phi P \frac{c^2}{c^2 + B^2} + \phi \left( L + Q \frac{c^2}{c^2 + A^2} w \right) (C - c)(1 + \delta) + \phi D \nabla^2 c - k_c c, \tag{SI15}$$

$$\frac{\partial w}{\partial t} = k_{i-}(1 - w) - k_{i+} w c. \tag{SI16}$$

Here, $c(x, y, t)$ is the cytosolic calcium concentration and $w(x, y, t)$ is the fraction of calcium release channels currently available. The factors $\phi$ multiplied with the pump, release, and diffusion terms ensure that these intracellular processes occur only within the cell. The additional decay term $-k_c c$ allows calcium to decay in regions that the cell has left.

As $w$ is a dimensionless gating variable, its equation is left unchanged from the tubule model and is not multiplied by $\phi$. Consequently, $w$ is solved over the entire domain, but is only physically meaningful within the cell interior, and extending it outside does not affect the cell dynamics.

#### SI4 Numerical implementation

The governing equations for the tubule model are solved numerically on a one-dimensional domain using a finite-difference discretisation with no-flux boundary conditions. Time integration is performed using the Euler–Maruyama scheme, with stochastic fluctuations applied to $Q(x, t)$.

The phase-field equations are solved numerically on a two-dimensional domain also using finite-difference methods with no-flux boundary conditions. The phase-field equations are solved numerically as stochastic partial differential equations, with fluctuations applied to the phase field and calcium variables, while the spatial profile of $Q(x, y)$ remains fixed.

Across all simulation setups, simulations are repeated 30 times to assess the robustness of the results. The full set of parameters is given by Table SI1.

| **Parameter** | **Value** | **Meaning** | **Model** |
|---|---|---|---|
| $R$ | $10^{-5}\,\mathrm{m}$ | Mean radius [4] | 1D |
| $d$ | $10^{-6}\,\mathrm{m}$ | Thickness of tubule wall [4] | 1D |
| $\mu$ | $2\,\mathrm{Pa\,s}$ | Viscosity of cytoplasm [4] | 1D |
| $\eta$ | $2^5\,\mathrm{Pa\,s}$ | Viscoelastic modulus of tubule wall [4] | 1D |
| $E_1$ | $5\times10^4\,\mathrm{Pa\,s}$ | Elastic constant of tubule wall [4] | 1D |
| $P_0$ | $1\times10^2\,\mathrm{Pa}$ | Theoretical maximum active pressure [28] | 1D |
| $\tau$ | $8\,N\,s\,m^{-2}$ | Time scale of membrane dynamics | Phase field |
| $\gamma$ | $1\times10^{-16}\,N$ | Passive surface tension | Phase field |
| $\epsilon$ | $1.8\times10^{-5}\,m$ | Total width of cell membrane and actin cortex | Phase field |
| $\beta$ | $1\times10^{-4}\,N\,m^{-2}$ | Volume constraint parameter for $\phi$ | Phase field |
| $\alpha$ | $2.5\times10^{-11}\,N\,m^{-1}$ | Active contractile tension | Phase field |
| $k_{hc}$ | $2.5\times10^{-1}\,s^{-1}$ | Spring like elastic parameter for height $h$ | Phase field |
| $k_{hv}$ | $2\,s^{-1}\,m^{-2}$ | Volume constraint parameter for $h$ | Phase field |
| $k_{h-}$ | $1\times10^{-2}\,s^{-1}$ | Decay parameter for $h$ | Phase field |
| $k_c$ | $1\times10^{-1}\,s^{-1}$ | Decay parameter for $c$ | Phase field |
| $k_1$ | $2\times10^{-2}\,\mathrm{s}^{-1}$ | Effective rate of actin polymerization [4] | Both |
| $k_2$ | $5\times10^{-3}\,\mathrm{s}^{-1}$ | Effective rate of actin depolymerization [4] | Both |
| $\bar{Q}$ | $0.3\,\mathrm{s}^{-1}$ | Mean maximum channel permeability [5] | Both |
| $L$ | $2.5\times10^{-3}\,\mathrm{s}^{-1}$ | Calcium leak rate from vacuoles [5] | Both |
| $k_{i+}$ | $2.5\times10^4\,\mathrm{mol\,s}^{-1}$ | Forward rate constant of $Ca^{2+}$ channel binding [5] | Both |
| $k_{i-}$ | $3.75\times10^{-3}\,\mathrm{mol\,s}^{-1}$ | Reverse rate constant of $Ca^{2+}$ channel binding [5] | Both |
| $A$ | $1.18\times10^{-7}\,\mathrm{M}$ | Phenomenological rate parameter [5] | Both |
| $B$ | $1.18\times10^{-7}\,\mathrm{M}$ | Phenomenological rate parameter [5] | Both |
| $P$ | $1.18\times10^{-7}\,\mathrm{M\,s}^{-1}$ | Phenomenological calcium pump parameter [5] | Both |
| $\delta$ | 19 | Morphological parameter [5] | Both |
| $D$ | $5\times10^{-10}\,\mathrm{m}^2\,\mathrm{s}^{-1}$ | Diffusion constant of $[Ca^{2+}]$ in cytoplasm [20] | 1D |
| $D$ | $1\times10^{-10}\,\mathrm{m}^2\,\mathrm{s}^{-1}$ | Diffusion constant of $[Ca^{2+}]$ in cytoplasm | Phase field |

Table SI1: Summary of model parameters, including their values, physical interpretations, corresponding references for the parameter values, and the model(s) in which they are used.

### SI5 Metrics

#### SI5.1 Tubule model

We use several metrics to characterize the oscillations of the tubule and to quantify how the spatiotemporal dynamics change following external stimulation. All metrics are computed using $N = 30$ independent simulations.

**Phase relationship analysis.** To quantify the phase relationship between calcium, pressure, and radius, we performed a phase-binned analysis at the tubule midpoint. For each simulation, the final $5,000\,\mathrm{s}$ of steady-state calcium dynamics were mean-centered and the instantaneous phase for the calcium oscillations was estimated using the Hilbert transform. The time points were then assigned to 32 phase bins, where the mean deviations of all variables were computed.

**Directional index.** To quantify the directionality of wave propagation, we define a directional index (DI) based on the asymmetry of the spatiotemporal Fourier spectrum. Let $s(x,t)$ denote the variable of interest (e.g., calcium concentration or radius). We compute its Fourier transform in space and time, $\hat{s}(k,\omega)$, and define the spectral energy as $E(k,\omega) = |\hat{s}(k,\omega)|^2$. The directional index is defined as

$$\mathrm{DI} = \frac{\sum_{k>0,\omega} E(k,\omega) - \sum_{k<0,\omega} E(k,\omega)}{\sum_{k>0,\omega} E(k,\omega) + \sum_{k<0,\omega} E(k,\omega)}, \tag{SI17}$$

where the sums are taken over the discrete Fourier modes. This measure ranges from $-1$ to 1, where positive values indicate a bias toward waves propagating in the positive spatial direction (in this case towards the food source), negative values indicate the opposite, and values near zero correspond to symmetric or non-directional dynamics.

**Dominant wavenumber.** To characterise the spatial structure of the oscillations, we compute the spatial Fourier spectrum of $s(x,t)$ and define the dominant wavenumber $k_{\mathrm{peak}}$ as the wavenumber at which the spectral power is maximal. This quantity reflects the characteristic spatial scale (wavelength) of the pattern.

**Left–right biomass asymmetry.** To quantify spatial biomass and calcium concentration asymmetry, we divide the domain into left and right halves and compute the time-averaged signal in each region. Denoting these by $L$ and $R$, respectively, we define the asymmetry as

$$\mathrm{Asymmetry} = \frac{L-R}{L+R}. \tag{SI18}$$

Positive values indicate higher concentration/biomass on the left side of the tubule, while negative values indicate higher concentration/biomass on the right side of the tubule.

**Changes before and after stimulation.** In each simulation, following an initial transient period of 10,000 s, each metric is computed over a 10,000 s window of steady-state dynamics before and after stimulation. The change in each metric (e.g., $\Delta$DI) is then computed as the difference between the values in the post-stimulation and pre-stimulation phases.

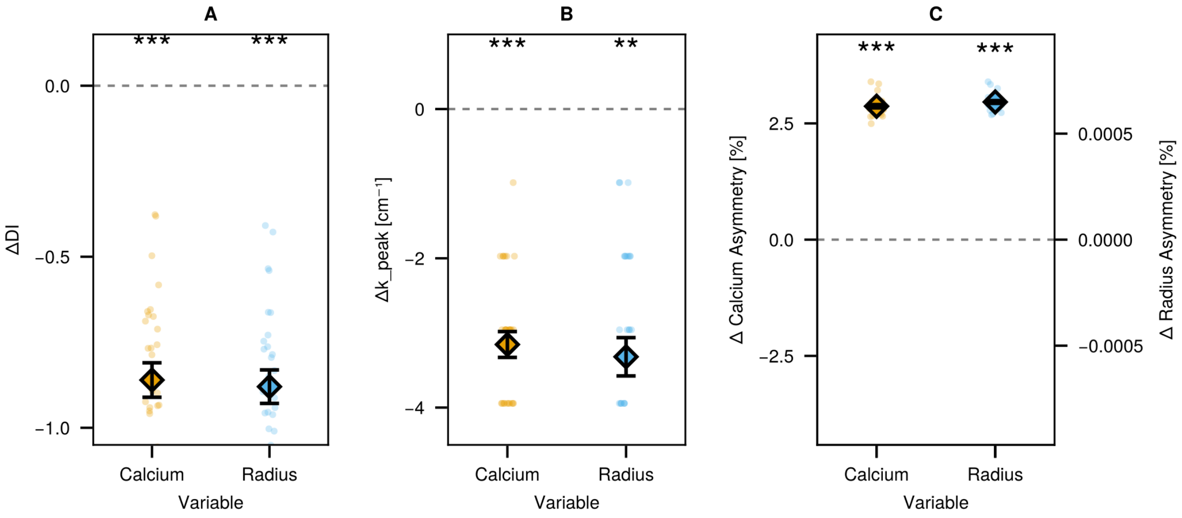


Fig. SI4: **Changes in spatiotemporal dynamics after introduction of a food source on the left side of the tubule.** Changes in directional index ($\Delta$DI), dominant wavenumber ($\Delta k_{\text{peak}}$), and spatial asymmetry following the introduction of asymmetric $Q$. For each simulation, metrics are computed over steady-state time windows before and after stimulation ($t = 10,000$), and differences were computed by taking the post- minus the pre-stimulation values. **(A)** Change in directional index for calcium and radius signals, showing a strong shift toward directional wave propagation toward the food source after stimulation. **(B)** Change in dominant wavenumber, indicating a decrease in $k_{\text{peak}}$, suggesting a shift toward larger spatial scales. **(C)** Changes in spatial asymmetry for calcium concentration (left axis) and radius (i.e. biomass) (right axis), showing redistribution toward the high-$Q$ side of the domain. Points represent individual simulation runs ($n = 30$), diamonds indicate the mean, and error bars denote the standard error of the mean. Horizontal dashed lines indicate zero change. Statistical significance is assessed using paired $t$-tests; $^{***}p < 0.001$, $^{**}p < 0.01$, $^{*}p < 0.05$.

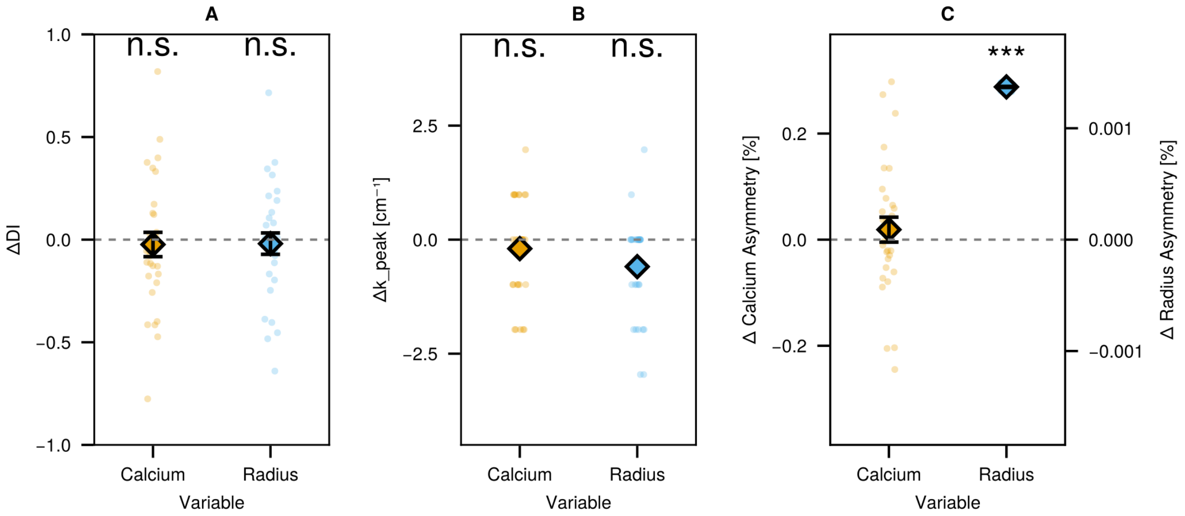


Fig. SI5: **Changes in spatiotemporal dynamics following local application of Cytochalasin D to the left side of the tubule.** Changes in directional index (ΔDI), dominant wavenumber ($\Delta k_{\text{peak}}$), and spatial asymmetry following local inhibition of actin polymerization, implemented as a local reduction in $k_1$. For each simulation, metrics are computed over steady-state time windows before and after stimulation ($t = 10,000$), and differences were computed by taking the post- minus the pre-stimulation values. **(A)** Change in directional index for calcium and radius signals, showing no significant change in wave directionality after stimulation. **(B)** Change in dominant wavenumber, indicating no significant change in the spatial structure of the oscillations. **(C)** Changes in spatial asymmetry for calcium concentration (left axis) and radius (right axis), showing a redistribution of radius (biomass) toward the perturbed side, while calcium asymmetry remains unchanged. Points represent individual simulation runs ($n = 30$), diamonds indicate the mean, and error bars denote the standard error of the mean. Horizontal dashed lines indicate zero change. Statistical significance is assessed using paired $t$-tests; ***$p < 0.001$, **$p < 0.01$, *$p < 0.05$.

#### SI5.2 Phase field model

To quantify spreading, migration, and food-source interaction in the phase-field simulations, we extract measures of cell shape, motion, and connectivity from the phase-field variable $\phi(x, y, t)$ and thickness field $h(x, y, t)$. The cell area is defined as the region where $\phi > 0.5$. As for the tubule model, each setup is simulated 30 times. Across all three setups (baseline, chemotaxis, and network formation), we compute the area covered by the cell over time. For the baseline experiment, we compute the front propagation speed. For the network formation experiment, we compute the number of connected food sources at each time point, to assess whether and when the cell splits. For the chemotaxis experiment, we compute the directional speed toward the food source, the leading-edge speed, and the center-of-mass speed, as well as the evolution of the leading-edge and center-of-mass positions over time.

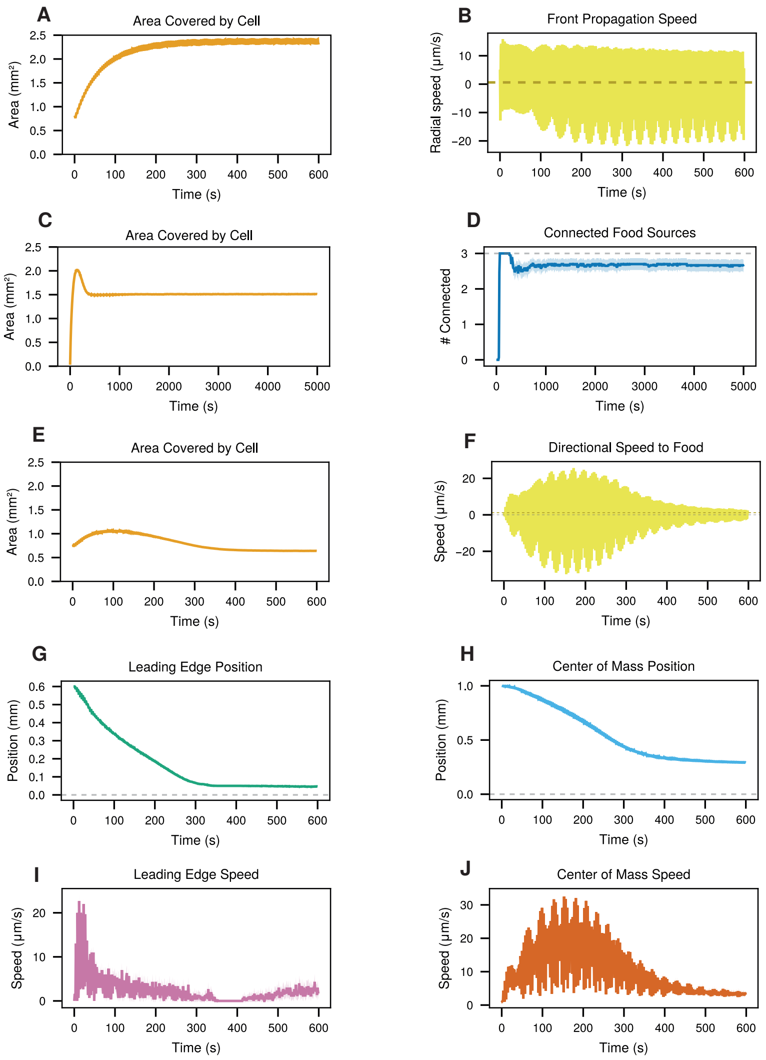


Fig. SI6: **Phase-field metrics across baseline, network formation, and chemotaxis experiments. (A,B)** Baseline experiment: cell area over time and front propagation speed, showing how the cell spreads until it covers most of the domain. The front propagation speed fluctuates between approximately $-20$ and $20\,\mu$m/s, with a net positive expansion. **(C,D)** Network formation experiment: cell area and number of connected food sources over time. The cell grows until it reaches the food sources and subsequently redistributes its biomass to form tubule-like connecting structures. In some simulations, the cell splits after reaching the food sources. **(E–J)** Chemotaxis experiment: cell area, directional speed toward the food source, leading-edge speed, and center-of-mass speed, together with the corresponding positions over time. The cell initially spreads radially and subsequently exhibits directed motion toward the food source. The directional speed fluctuates between approximately $-20$ and $20\,\mu$m/s, with a net positive bias toward the food. Lines indicate ensemble means and shaded regions indicate variability across simulations ($n = 30$).

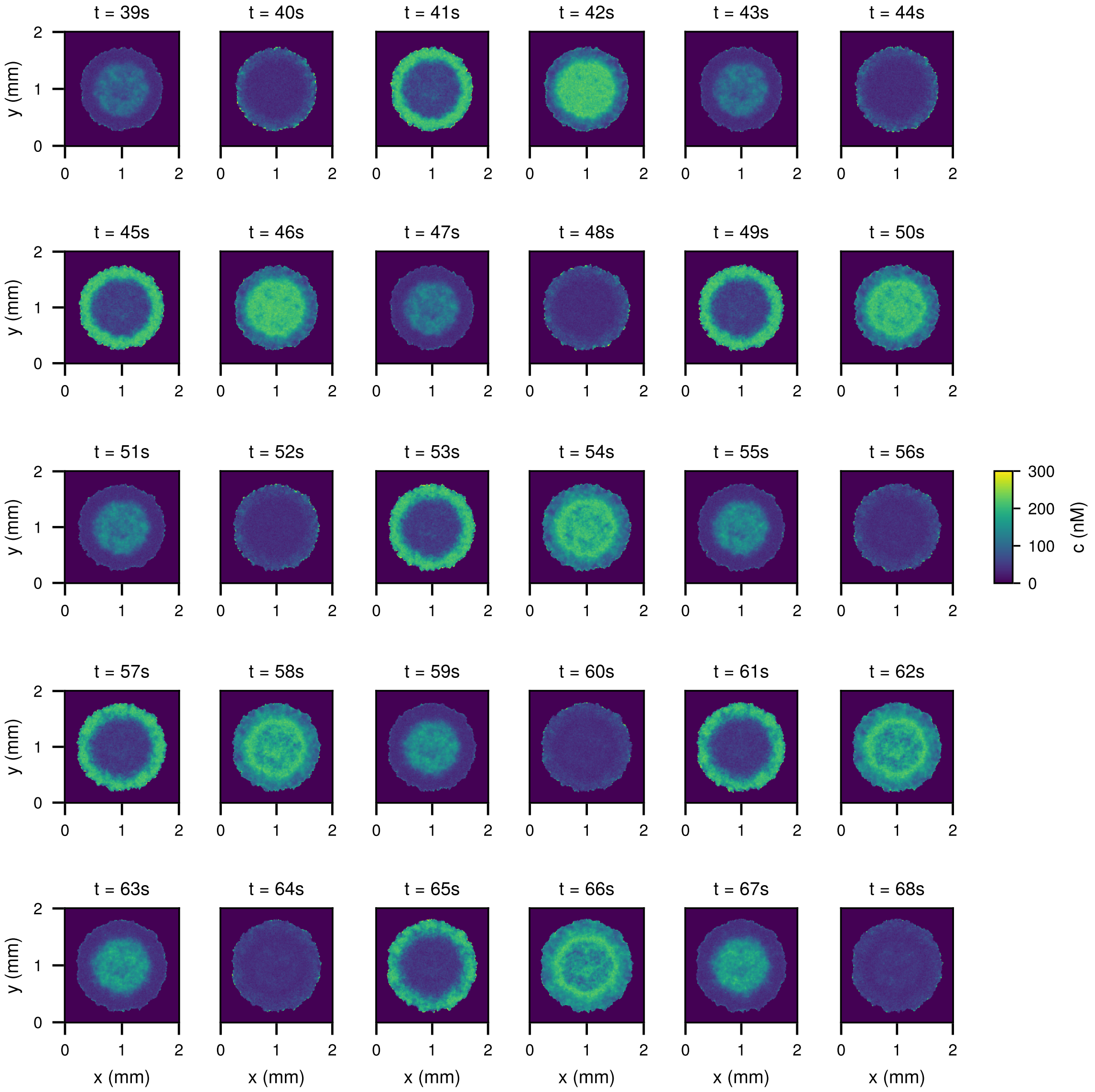


Fig. SI7: **Example time series of calcium dynamics in the absence of external cues.** Snapshots of the calcium concentration over 30 consecutive seconds from a representative simulation in the baseline experiment. The panels show oscillatory calcium dynamics with a radially symmetric spatial distribution that oscillates. These snapshots show the intrinsic calcium oscillations in the absence of external stimulation.

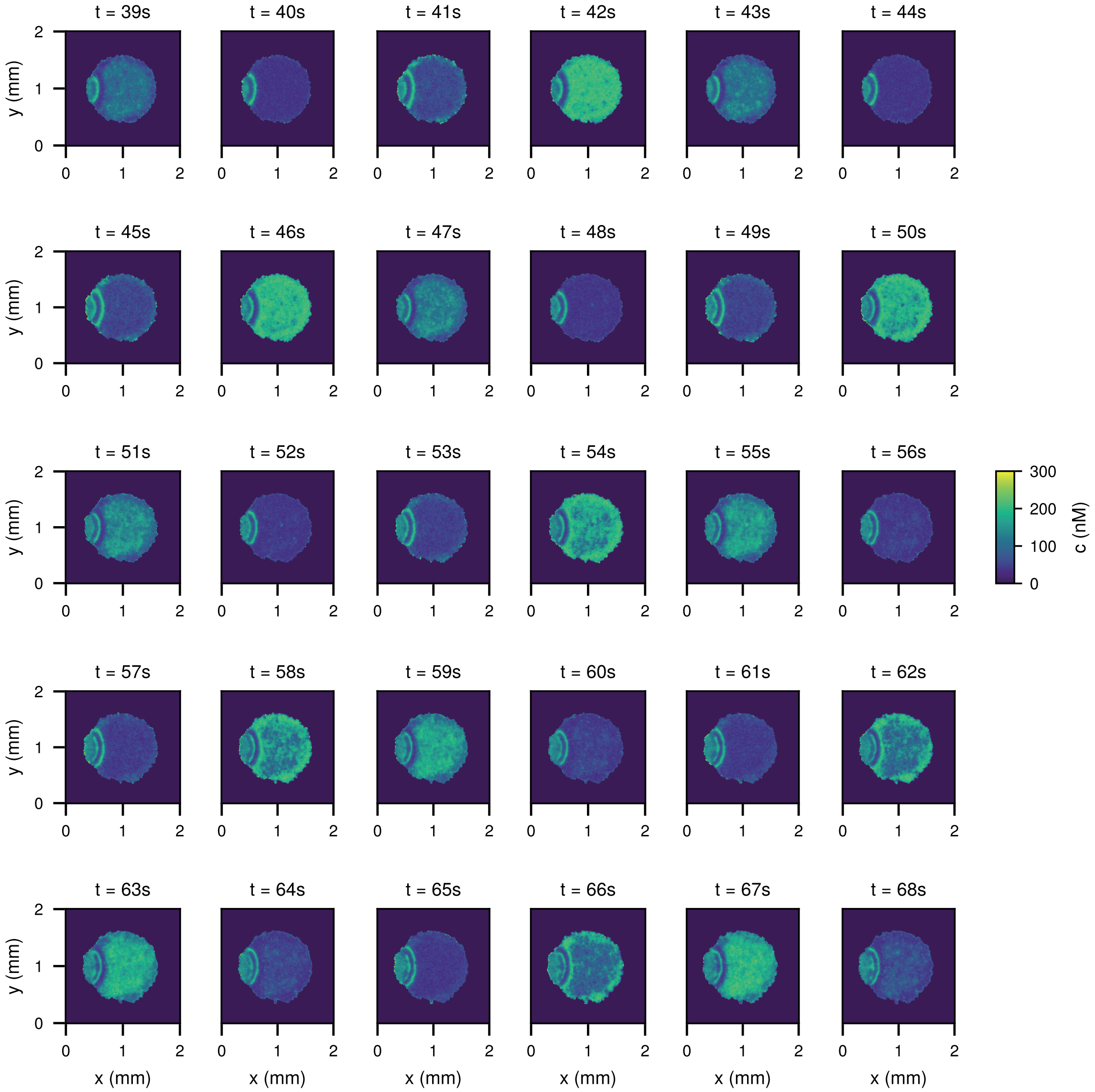


Fig. SI8: **Example time series of calcium dynamics during chemotaxis.** Snapshots of the calcium concentration over 30 consecutive seconds from a representative simulation in the chemotaxis experiment. The panels show oscillatory calcium dynamics as the cell comes into contact with the nutrient field, with oscillatory activity spreading and becoming spatially redistributed in response to the external cue.

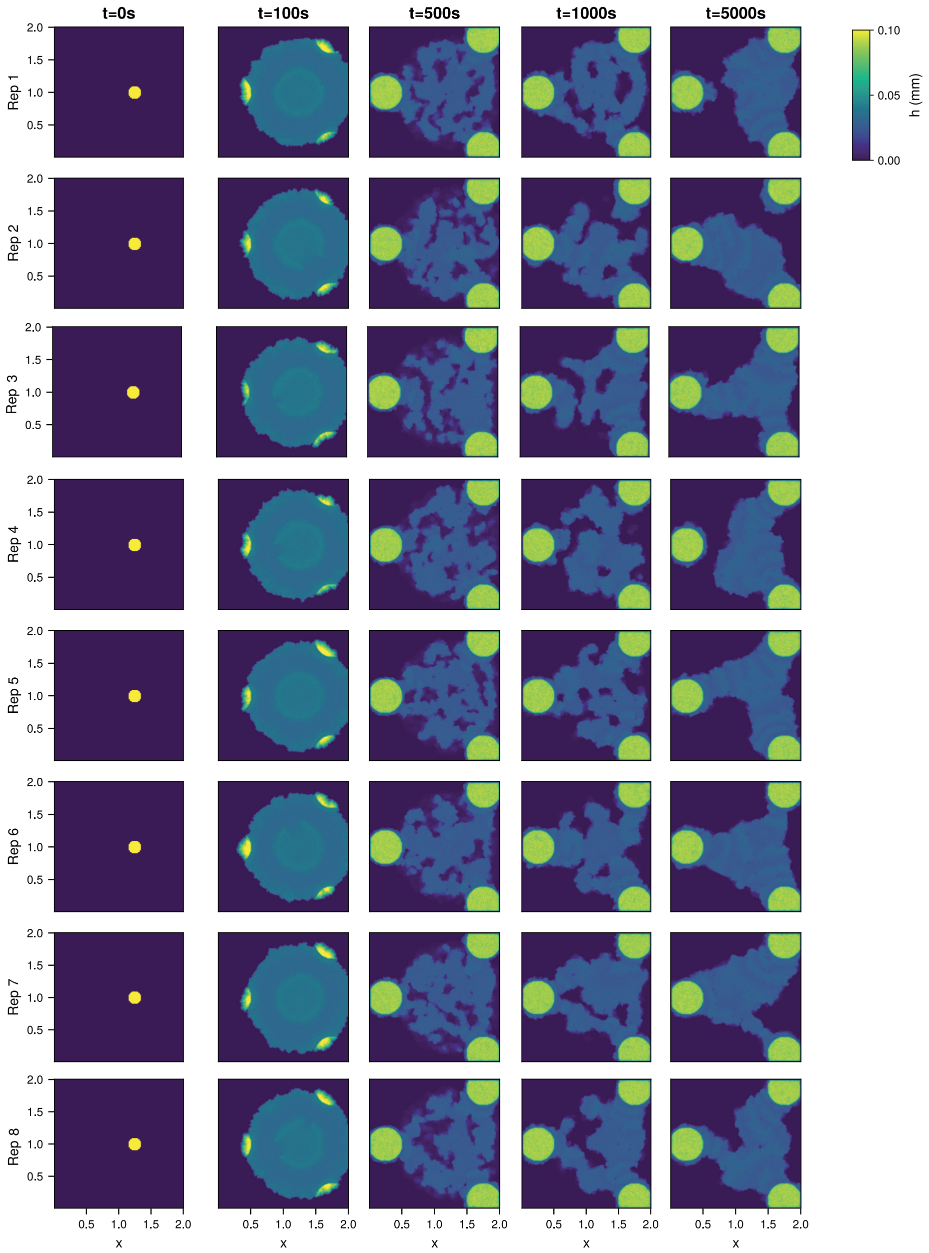


Fig. SI9: **Representative realizations of network formation dynamics.** Snapshots from multiple simulation runs of the network formation experiment at selected time points, with each row corresponding to a different replicate. The cell expands, connects to food sources, and forms network-like structures. In some realizations, the cell detaches upon contact with the food sources and splits into two separate cells.

### References


[1] K. Alim, G. Amselem, F. Peaudecerf, M. P. Brenner, and A. Pringle, "Random network peristalsis in physarum polycephalum organizes fluid flows across an individual," *Proceedings of the National Academy of Sciences*, vol. 110, no. 33, pp. 13306–13311, 2013.

[2] S. K. Ray, G. Valentini, P. Shah, A. Haque, C. R. Reid, G. F. Weber, and S. Garnier, "Information transfer during food choice in the slime mold physarum polycephalum," *Frontiers in Ecology and Evolution*, vol. 7, p. 67, 2019.

[3] E. B. Wilson, "Probable inference, the law of succession, and statistical inference," *Journal of the American Statistical Association*, vol. 22, no. 158, pp. 209–212, 1927.

[4] V. Teplov, Y. M. Romanovsky, and O. Latushkin, "A continuum model of contraction waves and protoplasm streaming in strands of physarum plasmodium," *Biosystems*, vol. 24, no. 4, pp. 269–289, 1991.

[5] J. Poledna, "Model of intracellular calcium oscillations activated by inositol trisphosphate," *General physiology and biophysics*, vol. 12, pp. 381–381, 1993.

[6] K.-E. Wohlfarth-Bottermann and M. Fleischer, "Cycling aggregation patterns of cytoplasmic f-actin coordinated with oscillating tension force generation," *Cell and Tissue Research*, vol. 165, no. 3, pp. 327–344, 1976.

[7] M. Ishigami, K. Kuroda, and S. Hatano, "Dynamic aspects of the contractile system in physarum plasmodium. iii. cyclic contraction-relaxation of the plasmodial fragment in accordance with the generation-degeneration of cytoplasmic actomyosin fibrils.," *The Journal of cell biology*, vol. 105, no. 1, pp. 381–386, 1987.

[8] H. Sugino and F. Matsumura, "Fragmin induces tension reduction of actomyosin threads in the presence of micromolar levels of ca2+.," *The Journal of cell biology*, vol. 96, no. 1, pp. 199–203, 1983.

[9] T. Hasegawa, S. Takahashi, H. Hayashi, and S. Hatano, "Fragmin: a calcium ion sensitive regulatory factor on the formation of actin filaments," *Biochemistry*, vol. 19, no. 12, pp. 2677–2683, 1980.

[10] Y. Yoshimoto and N. Kamiya, "Ca2+ oscillation in the homogenate of physarum plasmodium," *Protoplasma*, vol. 110, no. 1, pp. 63–65, 1982.

[11] R.-H. Tang, S. Han, H. Zheng, C. W. Cook, C. S. Choi, T. E. Woerner, R. B. Jackson, and Z.-M. Pei, "Coupling diurnal cytosolic ca2+ oscillations to the cas-ip3 pathway in arabidopsis," *Science*, vol. 315, no. 5817, pp. 1423–1426, 2007.

[12] G. J. Allen, S. P. Chu, C. L. Harrington, K. Schumacher, T. Hoffmann, Y. Y. Tang, E. Grill, and J. I. Schroeder, "A defined range of guard cell calcium oscillation parameters encodes stomatal movements," *Nature*, vol. 411, no. 6841, pp. 1053–1057, 2001.

[13] P. Launay, H. Cheng, S. Srivatsan, R. Penner, A. Fleig, and J.-P. Kinet, "Trpm4 regulates calcium oscillations after t cell activation," *Science*, vol. 306, no. 5700, pp. 1374–1377, 2004.

[14] M. J. Berridge, “The inositol trisphosphate/calcium signaling pathway in health and disease,” *Physiological reviews*, vol. 96, no. 4, pp. 1261–1296, 2016.

[15] J. Lévy, C. Bres, R. Geurts, B. Chalhoub, O. Kulikova, G. Duc, E.-P. Journet, J.-M. Ané, E. Lauber, T. Bisseling, *et al.*, “A putative ca2+ and calmodulin-dependent protein kinase required for bacterial and fungal symbioses,” *Science*, vol. 303, no. 5662, pp. 1361–1364, 2004.

[16] P. Schaap, I. Barrantes, P. Minx, N. Sasaki, R. W. Anderson, M. Bénard, K. K. Biggar, N. E. Buchler, R. Bundschuh, X. Chen, *et al.*, “The physarum polycephalum genome reveals extensive use of prokaryotic two-component and metazoan-type tyrosine kinase signaling,” *Genome Biology and Evolution*, vol. 8, no. 1, pp. 109–125, 2016.

[17] N. Matveeva, V. Teplov, and S. Beylina, “Suppression of the autooscillatory contractile activity of physarum polycephalum plasmodium by the inhibitor of the ip3-induced ca2+ release, 2-aminoethoxydiphenyl borate,” *Biochemistry (Moscow) Supplement Series A: Membrane and Cell Biology*, vol. 4, no. 1, pp. 70–76, 2010.

[18] A. Kochegarov, S. Beylina, N. Matveeva, G. Leontieva, and V. Zinchenko, “Ionomycin and 2, 5-di (tertbutyl)-1, 4,-benzohydroquinone elicit ca2+-induced ca2+ release from intracellular pools in physarum polycephalum,” *Comparative Biochemistry and Physiology Part A: Molecular & Integrative Physiology*, vol. 128, no. 2, pp. 279–288, 2001.

[19] T. Kato and Y. Tonomura, “Uptake of calcium ions into microsomes isolated from physarum polycephalum,” *The Journal of Biochemistry*, vol. 81, no. 1, pp. 207–213, 1977.

[20] B. S. Donahue and R. Abercrombie, “Free diffusion coefficient of ionic calcium in cytoplasm,” *Cell calcium*, vol. 8, no. 6, pp. 437–448, 1987.

[21] K. Natsume, Y. Miyake, M. Yano, and H. Shimizu, “Development of spatio-temporal pattern of ca2+ on the chemotactic behaviour of physarum plasmodium,” *Protoplasma*, vol. 166, no. 1, pp. 55–60, 1992.

[22] S. Bernhardt, M. Naim, U. Zehavi, and B. Lindemann, “Changes in ip3 and cytosolic ca2+ in response to sugars and non-sugar sweeteners in transduction of sweet taste in the rat.,” *The Journal of Physiology*, vol. 490, no. 2, pp. 325–336, 1996.

[23] M. Koganezawa and I. Shimada, “Inositol 1, 4, 5-trisphosphate transduction cascade in taste reception of the fleshfly, boettcherisca peregrina,” *Journal of neurobiology*, vol. 51, no. 1, pp. 66–83, 2002.

[24] L.-Q. Chen, “Phase-field models for microstructure evolution,” *Annual review of materials research*, vol. 32, no. 1, pp. 113–140, 2002.

[25] A. Moure and H. Gomez, “Phase-field modeling of individual and collective cell migration,” *Archives of Computational Methods in Engineering*, vol. 28, no. 2, pp. 311–344, 2021.

[26] D. Shao, W.-J. Rappel, and H. Levine, “Computational model for cell morphodynamics,” *Physical review letters*, vol. 105, no. 10, p. 108104, 2010.

[27] S. Alonso, M. Stange, and C. Beta, "Modeling random crawling, membrane deformation and intracellular polarity of motile amoeboid cells," *PloS one*, vol. 13, no. 8, p. e0201977, 2018.

[28] J.-P. Rieu, H. Delanoë-Ayari, S. Takagi, Y. Tanaka, and T. Nakagaki, "Periodic traction in migrating large amoeba of physarum polycephalum," *Journal of The Royal Society Interface*, vol. 12, no. 106, p. 20150099, 2015.
